# Genomics resolves historical uncertainties on phylogenetics and accommodates the systematics of marmosets and Goeldi’s monkey (Primates: Platyrrhini)

**DOI:** 10.1101/2023.06.10.544470

**Authors:** Rodrigo Costa-Araújo, Christian Roos, Fabio Röhe, José de Sousa e Silva, Patricia Domingues de Freitas, Alcides Pissinatti, Jean P. Boubli, Izeni P. Farias, Tomas Hrbek

## Abstract

Marmosets, with a total of 24 species classified into four genera (*Callithrix*, *Cebuella*, *Mico* and *Callibella*), are the smallest of the anthropoids and one of the most diverse and widespread groups of primates in South America. In contrast, the Goeldi’s monkey (*Callimico goeldii*) is represented by a single species of black, small, fungi-eating primates, endemic to west Amazonia. The phylogenetic relationships of marmoset genera and the phylogenetic position of Goeldi’s monkey, and consequently their systematics, remain uncertain and subject to debate because earlier studies revealed incongruent conclusions. Here we tackle this issue by first reviewing the systematics and the history of phylogenetic studies of marmosets and Goeldi’s monkey. We then explore their phylogenetic relationships by reconstructing a time-calibrated phylogeny using a genome-wide sampling of all lineages of marmosets, tamarins, Goeldi’s monkey, lion tamarins, capuchins, and squirrel monkeys. Our results clearly demonstrate that historical disagreements on phylogenetics and systematics of marmosets are due to incomplete lineage sorting, low phylogenetic signal of morphological and ecological characters, and low sampling at the DNA level. We show that Goeldi’s monkey is a sister lineage to marmosets and suggest that past incongruencies between studies on its phylogenetics and systematics are due to homoplasy of morphological characters traditionally used to infer primate relationships. Accordingly, we accommodate a genus-level classification for marmosets based on a fully-resolved phylogeny and multiple biological traits, redefine the genus *Mico*, update the definitions of *Callibella*, *Callithrix*, and *Cebuella*, and sediment the family-level classification of Goeldi’s monkey.

## Introduction

Marmosets are small monkeys that feed largely on tree exudates, have clawed-like nails, long incisors and short canines on the lower jaw (Coimbra-Filho & Mittermeier, 1976; Hershkovitz, 1977; Rosenberger, 2020). With 24 species in four genera (*Callithrix, Cebuella, Callibella, Mico*) distributed across South America, they form one of the most diverse and widespread groups among Platyrrhini (Figure 1; Boubli et al, 2021; Costa-Araújo et al, 2019, 2021; Culot et al, 2019; Rylands & Mittermeier, 2013). In contrast, the Goeldi’s monkey is represented by a single species, *Callimico goeldii* (Thomas, 1904), which is endemic to western Amazonia (Rylands & Mittermeier, 2013). This species relies largely on fungi in its diet and is small-bodied, has clawed-like nails, long incisors and short canines as marmosets, as well as three pairs of molars and the females give birth to singletons as capuchin and squirrel monkeys (Hershkovitz, 1977; Rosenberger, 2020).

**Figure 1.**
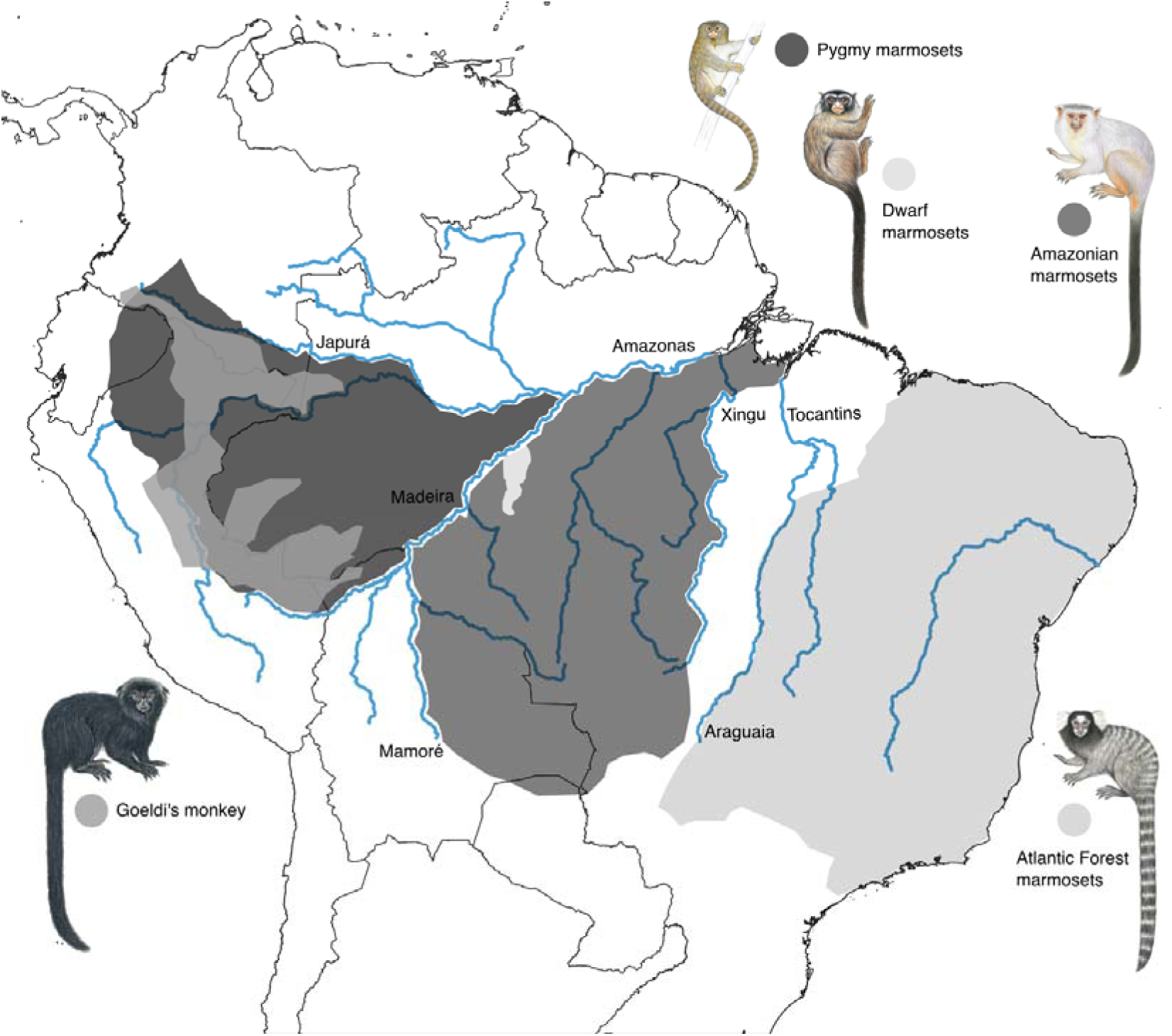
The geographic distribution of marmosets and Goeldi’s monkey in South America: the Atlantic Forest marmosets (e.g. *Callithrix jacchus*) in the east of Brazil (Culot et al, 2019), the Amazon marmosets (e.g. *Mico schneideri*) in southern Amazonia (IUCN, 2021), the dwarf marmoset (*Callibella humilis*) in southcentral Amazonia (Silva et al, 2018), the pygmy marmosets (e.g. *Cebuella pygmaea*) in northwest Amazonia (Boubli et al, 2018), and the Goeldi’s monkey (*Callimico goeldii*) in western Amazonia (IUCN, 2021). Illustrations: Stephen Nash.

Several hypotheses on the relationships and systematics of the marmosets and Goeldi’s monkey have been proposed in the past two and a half centuries, from Linnaeus (1758) to Rosenberger (2020). The majority of these derive from studies focusing on the morphological relatedness or phylogenetic relationships of platyrrhines (Primates, Platyrrhini; e.g. Elliot, 1913; Hershkovitz, 1977; Rosenberger, 1980; St.-Hilaire, 1812; Schneider et al, 1993, 2012; Simpson, 1945; Wang et al, 2018), and only a few of them focused specifically on marmosets (e.g. Ford & Davis, 1992; Natori, 1986; Sena et al, 2002; Tagliaro et al, 1997) and on the Goeldi’s monkey (e.g. Horovitz et al, 1998; Pastorini et al, 1998). Consequently, the hypotheses on evolutionary relationships and systematics of this group of primates are based on a reduced representation of specimens, species, a single source of data, and on a myriad of assumptions regarding the composition of taxonomic ranks made prior data collection and analysis.

In this sense, it is not surprising that the hypotheses on evolutionary relationships and systematics of marmosets and Goeldi’s monkey are extensively incongruent. Such incongruencies are observed when comparing studies based on the same or on distinct morphological or molecular characters, on studies with morphological versus molecular characters, or on descriptive versus phylogenetic approaches (Supplementary Material 1). To date, there are four schemes in use for the genus-level classification of marmosets (Table 1) and the Goeldi’s monkey has been classified in the Callitrichidae family, in its own family, Callimiconidae, or as a subfamily in Callitrichidae or Cebidae.

The relationships among marmosets and the phylogenetic position of *Cl. goeldii* remain unsettled because there is no clue on why different studies obtained results that are incongruent. Hypothesizing and using classifications for these taxa have largely followed conceptual or operational preferences, for example, which type of data and analysis generates “more reliable inferences” or which is “the correct” amount or kind of dissimilarity that grants a biological group a name and a given rank (e.g. Garbino et al, 2019). Therefore, classifications remain debated and the hypotheses have been accepted––or not––according to subjective criteria rather than to biological reasoning.

This means that, although rarely acknowledged, there are open questions regarding the phylogenetics and systematics of marmosets and the Goeldi’s monkey underlying the shallow debate of such conceptual and operational preferences. Why Amazon marmosets (*Mico*) have been recovered as sister taxa to pygmy (*Cebuella*) or to Atlantic Forest marmosets (*Callithrix*), depending on the source of data considered in the analysis? Why did different types of molecular markers recover distinct relationships between marmosets? Can morphological characters traditionally used in primate systematics inform evolutionary relationships of marmosets and the Goeldi’s monkey? Is the Goeldi’s monkey closely related to marmosets, to tamarins and lion tamarins, or is it the first offshoot among them? The answers to those questions would advance our knowledge on the evolutionary history of marmosets and the Goeldi’s monkey and support a stable systematic arrangement for this group of primates.

Here we address those questions firstly by reviewing the systematics and nomenclature of marmosets and the Goeldi’s monkey, and the hypotheses proposed to date on their phylogenetic relationships (Supplementary Material 1). We then explore the evolutionary relationships of these taxa reconstructing a time-calibrated phylogeny using a genome-wide sampling of marmosets, the Goeldi’s monkey, tamarins (*Leontocebus, Saguinus*, *Tamarinus*), lion tamarins (*Leontopithecus*), capuchins (*Cebus, Sapajus*), squirrel monkeys (*Saimiri*), and night monkeys (*Aotus*). Our results resolve the phylogenetic relationships among marmosets, the phylogenetic position of the Goeldi’s monkey, and reveal the causes of historical disagreements on the phylogenetic systematics of these monkeys.

Accordingly, we accommodate key biological data available on marmosets in a monophyletic, simple and meaningful classification, redescribe the genus *Mico* Lesson, 1812, provide updated definitions of the other names here suggested for the genus-level classification of marmosets, and sediment the phylogenetic position and the family-level classification of the Goeldi’s monkey.

## Methods

### Sampling design

We sampled the genome of species in all lineages of marmosets, tamarins, lion tamarins, and the Goeldi’s monkey. In order to infer the relationships of Goeldi’s monkey to their living close relatives (see Perelman et al, 2011), and to root and time-calibrate our trees we also sampled the genome of species in all lineages of the related capuchins, squirrel monkeys, and night monkeys. More specifically, we included one species from each of the four infra-generic lineages of *Mico* (Costa-Araújo et al, 2019, 2021)––hereafter Amazon marmosets, the two species of *Cebuella* (Boubli et al, 2021)––hereafter pygmy marmosets, *Callibella humilis*––hereafter dwarf marmoset, and one species from each of the three lineages of *Callithrix* (Malukiewicz et al, 2017)––hereafter Atlantic Forest marmosets, to investigate the evolutionary relationships between marmosets. In order to investigate the phylogenetic position of *Calllimico* among its living close relatives (Perelman et al, 2011; Wang et al, 2018) and to root and time-calibrate our trees we sampled the genome of *Cl. goeldii*, six species representing three of the four genera of tamarins (Brcko et al, 2022), two of the four lion tamarin species, one species of each of the two genera of capuchins (*Cebus*, *Sapajus*; Lynch-Alfaro et al, 2012), *Saimiri cassiquiarensis*, and *Aotus trivirgatus.* In total, we sampled the genome of 40 specimens from 21 species of platyrrhines (Supplementary Table 1). All applicable international, national, and/or institutional guidelines for the care and use of animals were strictly followed. All animal sample collection protocols complied with the current laws of Brazil.

### Generation and analysis of genomic data

After DNA extraction from muscle samples following Doyle & Doyle (1990), we used a modified protocol of ddRAD sequencing optimized for the IonTorrent PGM that permits simultaneous digestion, ligation and barcoded adapter incorporation (https://github.com/legalLab/protocols-scripts). Sequencing reads were processed using the pyRAD pipeline (Eaton, 2014). During de novo assembly, we used a minimum coverage of 6x per locus, assembling all fragments in the 320–400 bp range. Nucleotides with PHRED scores < 30 were excluded, as well as loci with more than three low quality nucleotides. Following demultiplexing and extraction of loci using the above criteria (steps 1–2 of the pyRAD pipeline), we proceeded with clustering of alleles within loci, and of loci across individuals, and the generation of the datasets for analyses (pyRAD steps 3–7). We generated a dataset with all individuals where a locus was included if it was present in at least 75% of individuals, which was then subjected to PartitionFinder2 (Lanfear et al, 2017) to estimate the optimal number of partitions. Site models, clock models, and trees were unlinked for each partition, which were allowed to evolve under an uncorrelated log normal model with clock rates and standard deviations of clock rates estimated. We used the calibrated Yule species tree model, constraining the age of the root of the phylogeny at 19.95±1.96 million years ago (Mya; soft calibration prior derived from Perelman et al, 2011). We ran the MCMC for 10^9^ generations, collecting 5 x 10^4^ samples in STARBEAST2 to generate a Bayesian species tree. Finally, we generated a Maximum Likelihood partitioned phylogenetic tree in RAxML-ng (Kozlov et al, 2019) using the same parameters as in the STARBEAST2 analysis, and a dataset which included one individual per species and loci present in at least 50% of the individuals in order to compare topologies generated with different inference methods.

## Results

The Bayesian (Figure 2) and Maximum Likelihood (Supplementary Figure 1) phylogenies with data partitioned recovered the same topology with nodes supported by posterior probabilities of 1.0 and non-parametric bootstrap values of 100%. The initial split occurred between Callitrichidae and Aotidae + Cebidae, 19.95 Mya. Aotidae represents a sister lineage to Cebidae and separated from it 17.29 Mya. Within Cebidae, *Saimiri* separated from capuchin monkeys 15.98 Mya, with a further split of *Cebus* and *Sapajus* 2.11 Mya. In Callitrichidae, the tamarin clade (*Leontocebus, Saguinus, Tamarinus*) split off first, 14.32 Mya, followed by lion tamarins (*Leontopithecus*) branching at 13.30 Mya. Within tamarins, *Leontocebus* branched off 5.38 Mya, followed by the split between *Saguinus* and *Tamarinus* 4.72 Mya. The Goeldi’s monkey (*Callimico*) forms a sister clade to marmosets and separated from them 11.24 Mya. Within the marmoset clade, *Callithrix* branched off at 3.77 Mya, followed by *Cebuella* at 3.41 Mya, and the split between *Callibella* and *Mico* occurred 1.75 Mya. Species cladogenesis in Callitrichidae occurred 0.20-0.63 Mya (*Mico*), 0.23 Mya (*Leontopithecus*), 0.47-1.62 Mya (*Callithrix*) and 1.45 Mya (*Cebuella*).

**Figure 2.**
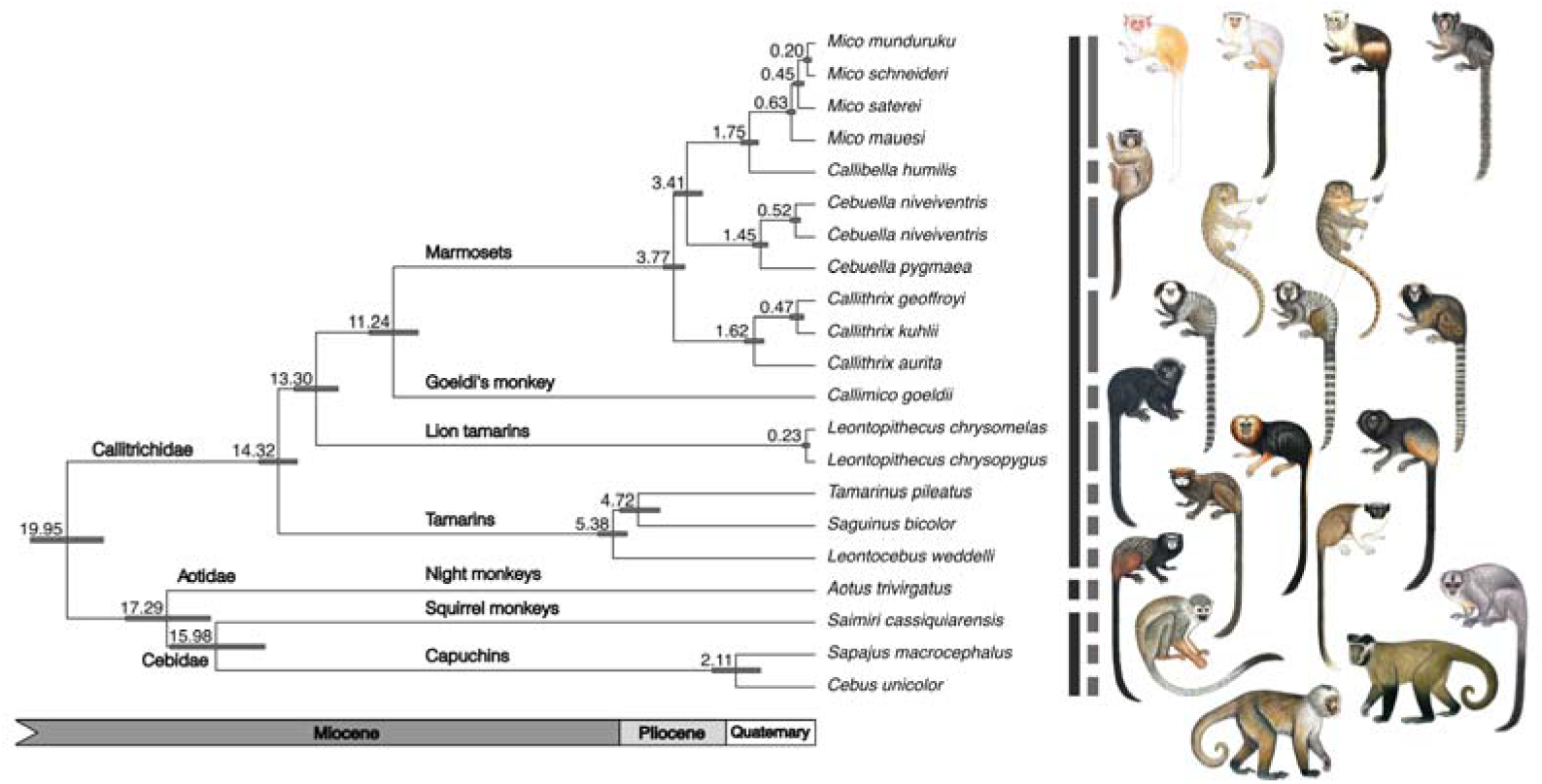
Time-calibrated phylogeny of Callitrichidae and outgroup families Cebidae and Aotidae reconstructed with Bayesian inference in BEAST2. Posterior probabilities are maximum at all nodes (pp=1.0). Number at nodes refer to time of divergence in million years and bars indicate 95% highest posterior densities. Illustrations: Stephen Nash.

The species tree generated with STARBEAST2 (Figure 3) revealed a highly consistent branching pattern among most taxa. However, for marmosets, three different evolutionary histories were recovered due to incomplete lineage sorting and retention of ancestral polymorphism. While the tree topology (((*Callibella*, *Mico*), *Cebuella*), *Callithrix*) (see also Figure 2) is supported by 84.75% of the loci sampled, 12.92% of the loci retrieved an unresolved branching pattern between *Callithrix*, *Cebuella* and *Mico*+*Callibella*, and 2.33% of the loci suggest *Cebuella* as sister clade to *Callithrix* and not to the *Mico*+*Callibella* clade due to incomplete lineage sorting and retention of ancestral polymorphisms.

**Figure 3.**
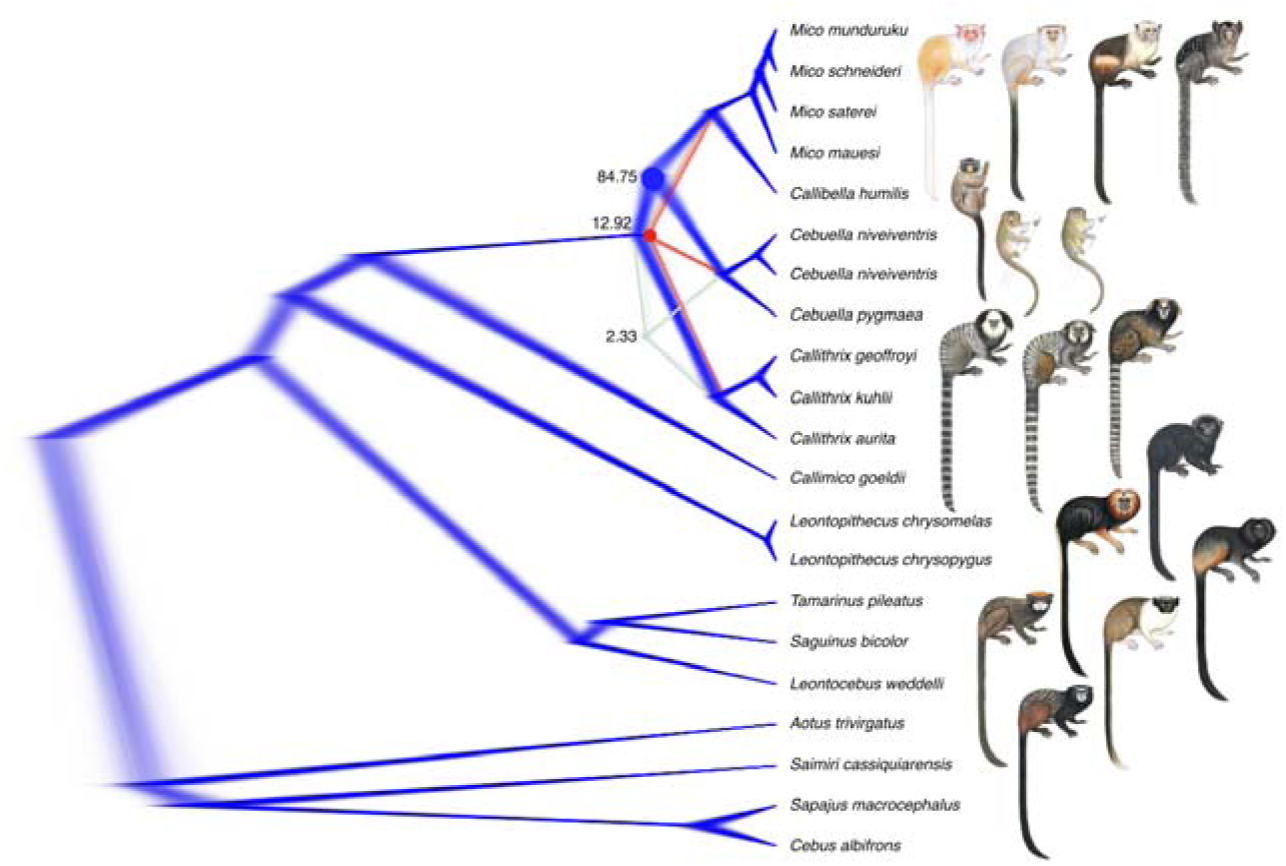
Phylogeny of Callitrichidae and outgroup families Cebidae and Aotidae reconstructed with STARBEAST2 using ddRAD data. Each colour line represents a topology according to different sets of loci. Proportion of loci/trees supporting divergent reconstructions are given at nodes, which are otherwise fully convergent.

## Discussion

### The causes of centenary incongruencies in marmoset phylogenetics and systematics

The first phylogenomic assessment of the evolutionary history of marmosets, presented here, clearly shows that incomplete lineage sorting due to retention of ancestral polymorphisms is the biological reason underlying historical incongruencies in the hypotheses of phylogenetic relationships and systematics of this group of primates. Our phylogenomic trees show a relatively short and overlapping interval between the divergence time of the Atlantic Forest marmoset clade (3.77 Mya) and the Amazon marmosets clade, and an elevated frequency of ancestral polymorphism retention (12.92%) by the Amazon + dwarf marmoset clade. The low frequency of ancestral polymorphism retention by the pygmy marmoset clade (2.33%) is best explained by a reduction in their effective population size at the time of divergence.

Due to incomplete lineage sorting three different topologies are obtained for marmosets, depending on which region of the genome is sampled––as it indeed has been shown in past phylogenetic analyses (e.g. Barroso et al, 1997; Meireles et al, 1998; Schneider et al, 2012; Sena et al, 2002; Tagliaro et al, 1997). In this sense, the low sampling coverage of the marmoset genome can be considered as an additional, operational issue underlying the lack of congruency in previous molecular phylogenetic hypotheses and their resulting systematic arrangements. That is, previous studies on marmosets recovered the relationships of a particular locus or loci––whether they shown different topologies or, by chance, a topology that coincides with the phylogenomic relationships presented here.

The retention of ancestral polymorphisms is also reflected in the morphology of marmosets and therefore in many previous phylogenetic and systematic hypotheses based on such source of information. *Callithrix* and *Mico* share a considerable proportion of alleles and have been long regarded as representatives of the same morphological group and genus (Hershkovitz, 1977; Rosenberger, 2020; St.-Hilaire, 1812; Spix, 1823) although they do not nest in a clade, and are not sister clades as shown here. *Callithrix* and *Cebuella* are also not sister taxa but as they share a lower proportion of alleles, as shown here, have been consistently and correctly considered as belonging to distinct morphological groups and genera (Cruz-Lima, 1944; Gray, 1870; Hershkovitz, 1977; Hill, 1957; Rosenberger, 2020; Rylands &Mittermeier, 2009; Rylands et al, 2009, 2012; Thomas, 1922). The retention of ancestral polymorphisms has, therefore, mislead morphology-based classifications of marmoset genera because the phylogenetic signal of the morphological characters traditionally adopted in such studies is inadequate to infer their evolutionary relationships.

Similarly, ecological traits in general do not bear enough phylogenetic signal that could allow an inference of the evolutionary relationships in marmosets. According to available information (see Rosenberger, 2020), *Callithrix* and *Mico* species can be considered as ecological equivalents in different biomes, they share a high number of loci, are morphologically similar but are not sister taxa. On the other hand, *Callibella* and *Mico* are sister taxa but occur in sympatry in Amazonia (Costa-Araújo et al, 2020; Silva et al, 2018), which is a clear evidence of niche shift and ecological differentiation in theoretical grounds (Carscadden et al, 2020). In fact, there are several examples of syntopic but ecologically distinct species classified in separate genera in primates (Lynch-Alfaro et al, 2012), felids (Di Bitetti et al, 2010; Sollmann et al, 2012), canids (Jácomo et al, 2004; Vieira & Port, 2007), otters (Moraes et al, 2021) and perissodactyls (Desbiez et al, 2009). Ecological evidence could shed light only on the phylogenetic relationship of *Cebuella* among marmosets, as the two species of *Cebuella* are ecologically distinct from *Mico* and *Callithrix* (Soini 1993; Hershkovitz, 1997; Rosenberger, 2020).

### The genus-level classification of marmosets

Considering the radiation of marmosets into four monophyletic lineages and the well-known morphological disparity between these lineages (Supplementary Material), we consider three plausible options for the marmosets’ classification at the genus level. These three classification schemes vary in the amount of information that can be retrieved from each and in their complexity of nomenclature and ranking (Figure 4).

I. Four genera: each of the four monophyletic lineages receivesfull genus status; *Callibella* for dwarf marmosets, *Callithrix* for Atlantic Forest marmosets, *Cebuella* for pygmy marmosets, and *Mico* for Amazon marmosets (Figure 4–A).
II. One genus with four subgenera: the basal marmoset clade receives full genus status, *Callithrix*, and each lineage receives the subgenus status: *Callithrix* (*Callibella*), *Callithrix* (*Callithrix*), *Callithrix* (*Cebuella*), and *Callithrix* (*Mico*) (Figure 4–B).
III. One genus without subgenera: the basal marmoset clade receives full genus status, *Callithrix*, applicable to the four lineages therein (Figure 4–C).

**Figure 4.**
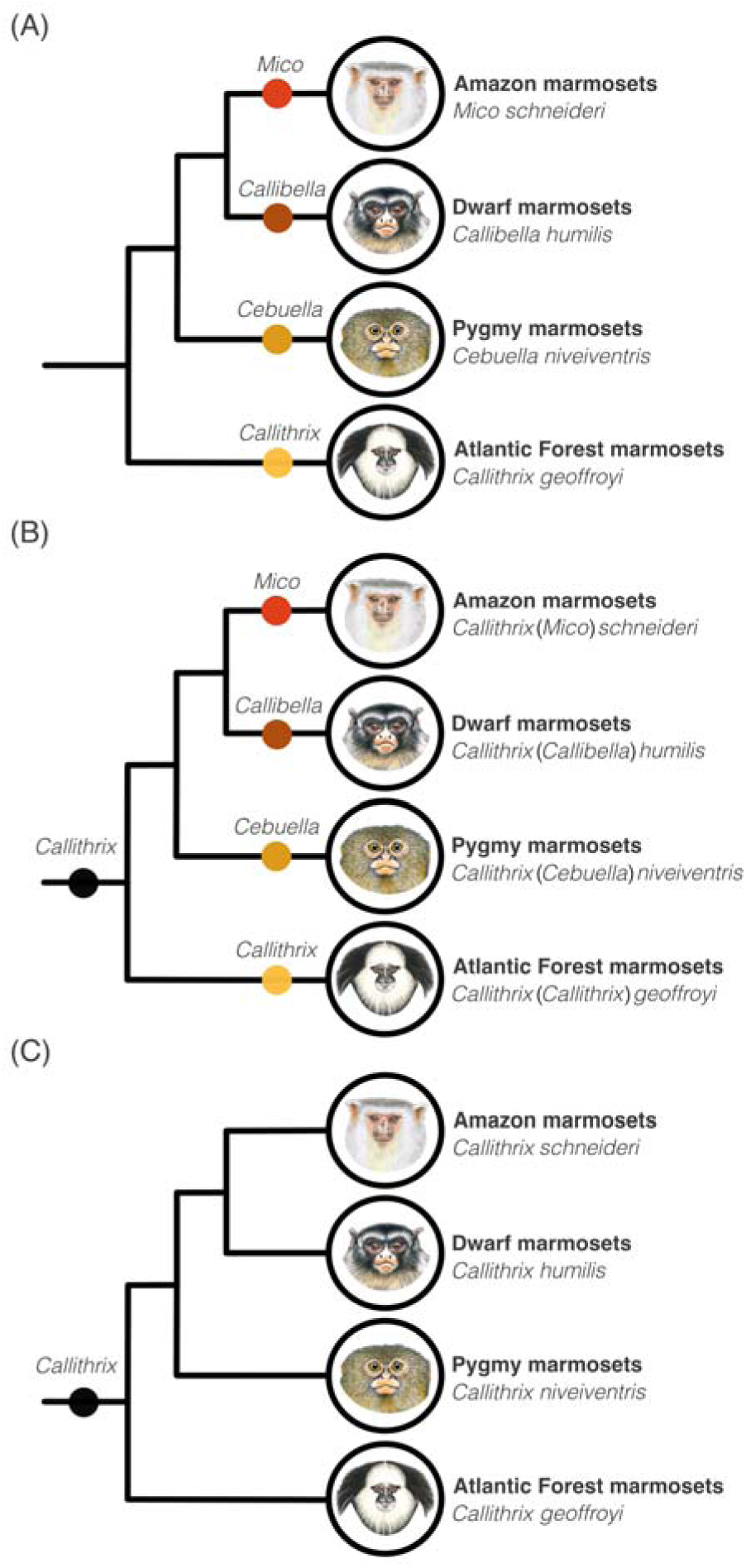
The three plausible options for the classification of the four marmoset lineages according to phylogenomic relationships (see figures 2, 3) and multiple biological traits (Supplementary Material). Illustrations: Stephen Nash.

The four genera scheme (I) follows thoroughly the phylogenomic relationships here presented by granting the full genus status to each of the four evolutionary lineages of marmosets; this option serves as a comprehensive system of information retrieval regarding the variation in body size, mass, skull, jaw and post-crania, pelage patterns, and zoogeography of each lineage (Supplementary Material). Moreover, it is a simple classification in terms of ranking and nomenclature. In fact, this scheme has been in use and it is widely accepted (e.g. IUCN, 2023; Rylands & Mittermeier, 2013), proving its meaningfulness for taxonomy users.

The four subgenera scheme (II) is informative and follows the phylogeny thoroughly as the four genera scheme but is the more complex and less practical option in terms of ranking and nomenclature. Compared to the four genera scheme it requires the additional ranking and naming of the basal marmoset clade and therefore an additional name and rank to properly designate genera and species. The single-genus scheme (III), although the simpler in ranking and nomenclature, does not retain the biological variation of the marmoset lineages nor follows phylogeny thoroughly.

The adoption of species groups within a single genus classification (e.g. Hershkovitz, 1977) is also plausible but adds unnecessary complexity with an informal rank and additional terminology. The use of three genera (Garbino, 2015; Schneider et al, 2012; Schneider & Sampaio, 2015), two genera (Rosenberger, 2020), or a subgenus for a single but not all the marmoset lineages (Garbino et al, 2019) is inadequate. These schemes do not mirror phylogeny––i.e. are not monophyletic classifications––and do not reflect the biological disparity of the four marmoset lineages.

We recommend the use of the four genera scheme to classify marmosets (Figure 5) because this is the simpler, correct option available in terms of ranking and nomenclature, serves as a robust system of information retrieval, is monophyletic, follows phylogeny thoroughly, and fulfils the goals of a classification (Vences et al, 2013). Such scheme (a) designates monophyletic lineages as genera, (b) retains information on key biological traits of each marmoset genus, and does not need (c) two levels of genus ranking and naming, (d) the superfluous classification of basal marmoset clade, nor the (e) adoption of species groups. In order to facilitate the use of such classification we re-describe the genus *Mico* and provide updated definitions for *Callibella*, *Callithrix*, and *Cebuella* (Supplementary Material).

**Figure 5.**
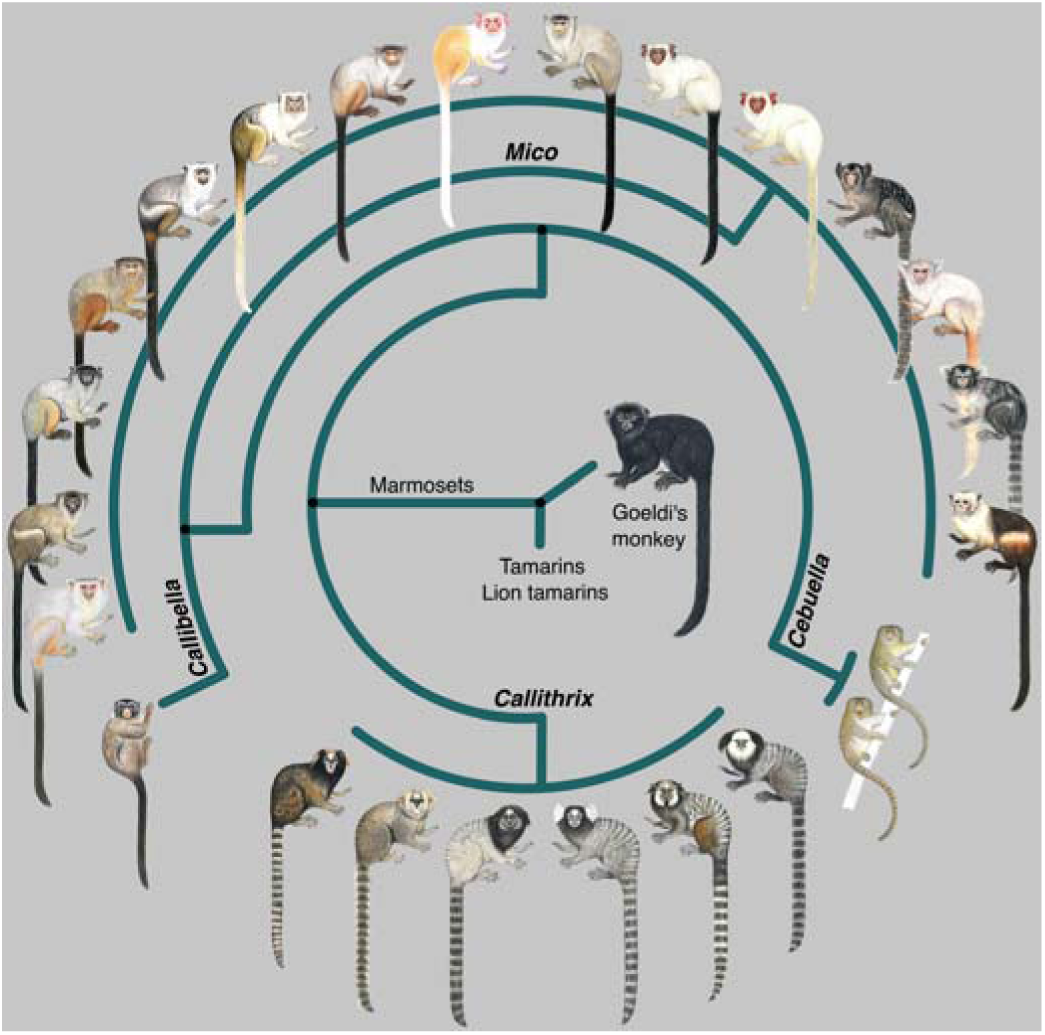
The marmosets and Goeldi’s monkey diversity (see Table 1), based on the phylogenomic inference presented here and multiple biological traits. Illustrations: Stephen Nash.

### Phylogenetics and classification of the Goeldi’s monkey

We unambiguously recovered the Goeldi’s monkey as sister taxon to the marmoset clade. This relationship has been retrieved by the majority of molecular phylogenetics studies (e.g. Osterholz et al, 2009; Pastorini et al, 1998; Perelman et al, 2011; Schneider et al, 1993; Wang et al, 2018; Wildman et al, 2009), whereas morphology-based inferences have retrieved *Callimico* as a sister taxon to *Leontopithecus* or as a first offshoot among callitrichids (e.g., Garbino, 2015; Rosenberger, 2020). Nonetheless, the taxa more closely related to *Cl. goeldii* are the extinct *Carlocebus* and *Patasola* (Horovitz, 1999). The incongruencies between past phylogenetic studies and classifications of *Cl. goeldii* are due to homoplasy of morphological characters used to infer its evolutionary relationships. Our results indicate that morphological characters adopted in previous studies generated phylogenies and classifications that mirror morphological affinities of homoplastic characters, not genealogy.

Therefore, there is no biological evidence for considering *Callimico* as an evolutionary intermediate between cebids and callitrichids or as the first offshoot among callitrichids. In turn, there is no reason for classifying Goeldi’s monkey in its own family or in a subfamily of Callitrichidae––unless other subfamilies or families are raised to classify marmosets, tamarins, and lion tamarins. Noteworthy, Hapalidae has priority over Callitrichidae (Groves, 2008) but Callitrichidae has been the most frequently used name since its proposition (Gray, 1821; Thomas, 1903b) and used ubiquitously in the last 50 years. Therefore, Hapalidae is de facto *nomen oblitum* and Callitrichidae (Figure 6) should be used for the family classification of *Callimico*.

**Figure 6.**
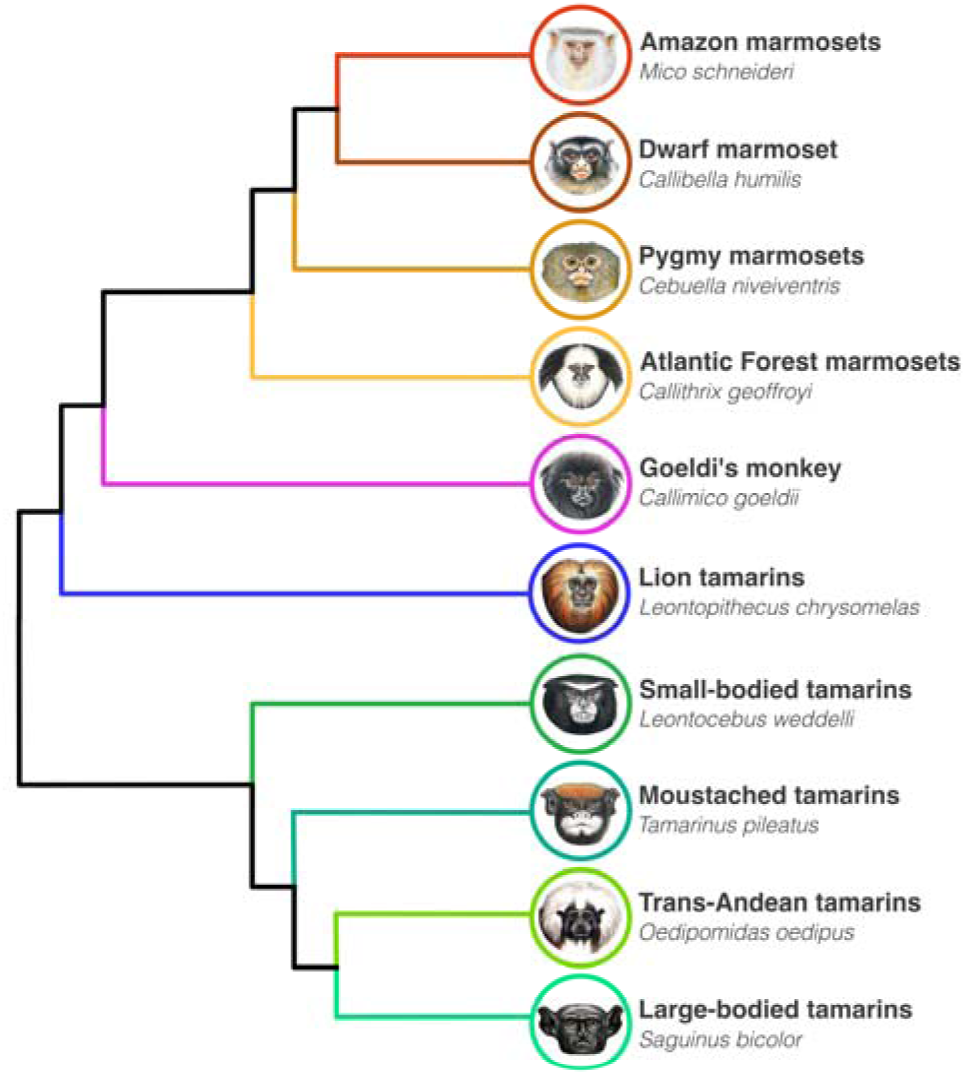
Phylogenetic relationships and genus-level classification of the extant lineages of Callitrichidae family (this paper, Brcko et al, 2022). Illustrations: Stephen Nash.

## Conclusion

Here we shown that the historical incongruencies on the evolutionary relationships and the instability of the genus-level systematics of marmosets are due to retention of ancestral polymorphisms, associated with the inadequateness of the morphological characters traditionally used in morphology-based phylogenies and the reduced sampling at the DNA level in molecule-based phylogenies. The retention of ancestral polymorphisms confounded previous morphology-based studies, which recovered the morphological relationships of these lineages but not necessarily their phylogenetic relationships. Due to the same reason, and also due to low sampling at the DNA level, previous molecular phylogenies recovered the phylogenetic relationships of the particular locus or loci analysed, which coincided or not with the history of the lineages. As shown here, three different topologies can be recovered depending on which portion of the genome is analysed.

We definitely demonstrate that the living sister taxa of the Goeldi’s monkey are the marmosets and that past uncertainties in its phylogenetic position and family-level classification are due to homoplasy of morphological characters used to infer its evolutionary relationships. In order to have a classification for marmosets that follows phylogeny thoroughly, accounts for their morphological disparity, and it is unnecessarily complex in ranking and nomenclature, the four genera classification scheme––*Callibella*, *Callithrix*, *Cebuella*, *Mico––*is the option. *Callimico* must be classified in the Callitrichidae family, as the sister taxon to marmosets.

Our results also highlight that the divergence time thresholds suggested by Goodman et al (1998) are not reliable proxies for morphological, ecological, and phylogenetic diversification in Callitrichidae and Cebidae. Therefore, these should not be used as a rule-of-thumb criterion in taxonomy and systematics of Platyrrhini. *Callibella* and *Mico* (1.75mya), *Ce. pygmaea* and *Ce. niveiventris* (1.45mya), and *Sapajus* and *Cebus* (2.11mya) all diverged from their closely related lineages earlier than suggested (4-6mya) to justify a classification in separate genera according to Goodman’s et al (1998) thresholds of divergence time. Divergence times must be revised to allow their use as proxies of diversification and as an accessory source of information in taxonomy and systematics of primates.

As taxonomy and systematics hypotheses increasingly include molecular data and adopt an integrative framework, changes in long-standing classifications and nomenclature based only on morphology data will become even more frequent. Many groups of Platyrrhini were not studied in the light of genomic data or subjected to proper molecular phylogenetic inferences (e.g. *Alouatta*, *Aotus*, *Ateles*, *Lagothrix*, *Pithecia*, *Saimiri*). Instead of “taxonomic inflation” or “instability”, the changes in Platyrrhini systematics so far reflect primarily the increase of our knowledge on the patterns of biological organization, evolution, and their underlying mechanisms. Such advances are rather welcome as they are paramount for science-based conservation and changes in human behaviour that are urgently needed to tackle ongoing biodiversity crisis, zoonotic pandemics, and climate change.

## Author contributions

R.C.A. and T.H. conceived the research, R.C.A, T.H., I.P.F., C.R., F.R., J.S.S.Jr., J.P.B, P.D.F. and A.P. provided materials, R.C.A., T.H. and I.P.F. generated genomic data, R.C.A. and T.H. analysed the data, R.C.A., T.H., C.R. and J.S.S.Jr. authored the drafts of the paper, R.C.A. prepared figures and tables, all authors reviewed the manuscript and approved the final version.

## Acknowledgements

This article is part of R.C.A.’s doctoral thesis carried out at the graduate program in Ecology of the National Institute of Amazon Researches, Manaus, Brazil, thanks to funding from Conselho Nacional de Desenvolvimento Científico e Tecnológico (CNPq; Grant Nos. 563348/2010, 140039/2018-1), Coordenação de Aperfeiçoamento de Pessoal de Nível Superior (CAPES; Grant Nos. 3261/2013, 001), Fundação de Amparo à Pesquisa do Amazonas (Grant No. 06200889/2019), Conservation Leadership Programme (Grant No. F02304217), Re:wild’s Margot Marsh Primate Action Fund (Grant No. 6002856), Idea Wild, National Science Foundation (Grant No. 1241066), Fundação de Amparo à Pesquisa de São Paulo (Grant No. 12/50260-6), and invaluable support from Brisa Araújo and Renata Sarmento. The article was further improved at the Mastozoology collection of the Goeldi’s Museum (Grant CNPq Nos. 316321/2020- 6, 300524/2021- 8, 301327/2021- 1, 302061/2021-5, 300936/2022- 2), Belém, Brazil, and at the Genetic department of the German Primate Center (Grant CAPES–Alexander von Humboldt stiftung No. 88881.512895/2020-01), Göttingen, Germany. R.C.A. is grateful to Raimundo Rodrigues, Ararão, Zagaia, Banha, Catitu, Dona Eunice, Marina e João Lutz, Natal, Seu Luís, Seu Roberto, Ozébio, Reginaldo, Canguru, Pamela Sateré-Mawé e Valdinelis, Angelisson e o povo Tenharin, Nilcelio e o povo Djahui, Rosangela e o povo Parintintin, aos amigos da TI Cachoeira Seca, Valdir e Jacaré, Fazenda ONF São Nicolau, Igor J. Roberto, Raony Alencar, Gustavo Canale, Fabiano Melo, several local families of Amazonia, and the teams of CPB and Itaituba bureau of ICMBio for making the collection of field data possible, as well as Fabricio Bertuol and Sandra Hernandez for the support on laboratory work, and Stephen Nash for sharing his art designs.

