## Supplementary material for "Genomics resolves historical uncertainties on phylogenetics and accommodates the systematics of marmosets and Goeldi’s monkey (Primates: Platyrrhini)"

**Supplementary Material. The systematic phylogenetic of marmosets and Goeldi's monkey**

*An overview of the history of classification and nomenclatural acts on marmosets and Goeldi's monkey*

Western scholars are aware of the Atlantic Forest marmosets since the 16th century as a result of the first invasions of European nations in South America, when these monkeys were painted as "new world novelties". The first marmoset painting known date back to 1541 (Hershkovitz, 1977), a few decades after the Portuguese ships arrived to Brazil. Only two centuries later the marmosets started to be studied scientifically, by Buffon & Daubenton (1756), whom described aspects of the external morphology and anatomy of internal organs of *Callithrix jacchus* Erxleben, 1777 and *Mico argentatus* (Linnaeus, 1771). These authors described marmoset and tamarin species, and set the seminal basis of their family systematics.

Buffon & Daubenton (1756) arranged the "Ouistiti" (*C. jacchus*), "Mico" (*M. argentatus*), "Pinche" (*Saguinus oedipus*), "Tamarin" (*S. midas*), "Marikina" (*Leontopithecus rosalia*), and "Saki" (*Pithecia pithecia*) in the "Sagoins", a group separate of the other Platyrrhini—the "New World Monkeys"—which is very similar to the Callitrichidae family (Rylands & Mittermeier, 2013). Linnaeus (1758) thereafter classified all the world's non-human primates in the genus *Simia* and it took a century of research to have marmosets and callitrichids back into a minimally informative scheme within the Linnean system. In fact, this is not an isolated case; the literature on morphological affinities and the systematics of marmosets and Goeldi's monkey is plenty of misconceptions. A myriad of names for the same taxa and incongruent classifications and phylogenetic hypotheses were proposed in the past 260 years for this group of monkeys.

Subsequently to Buffon & Daubenton (1756) and Linnaeus (1758), Erxleben (1777) included back all Sagoins in a separate group, the genus *Callithrix*, considering the lack of a prehensile tail, as Buffon & Daubenton (1756). *Callithrix* is the first genus name valid for marmosets, as pointed out by Hershkovitz (1977). Illiger (1811) improved such classification based on characters of dentition and tail by erecting the genus *Hapale* only for the Sagoins except *P. pithecia*. Illiger (1811) adopted *Callithrix* for *Cebus capucinus* and *Saimiri* cf. *sciureus*, and included both *Hapale* and *Callithrix* genera in the family Quadrumana. The only marmosets known to science until then were *C. jacchus* and *M. argentatus*.

St.-Hilaire (1812), based on the collections of Alexandre Rodrigues Ferreira's in Brazil—taken out of the Lisbon museum (Conselho Federal de Cultura, 1972)—created the genus *Jacchus* exclusively for marmosets relying on the number of molar teeth, non-prehensile tail, and claw-like nails. St.-Hilaire (1812) included in such genus *C. jacchus*, *M. argentatus*, plus two Amazonian marmosets (*M. humeralifer*, *M. melanurus*) and three Atlantic Forest marmosets (*C. aurita*, *C. geoffroyi*, *C. penicillata*) he described as new species. St.-Hilaire (1812) used *Callithrix* for squirrel (*Saimiri*) and titi monkeys (*Callicebus*, *Cheracebus*, *Plecturocebus*) as Illiger (1811) and created the family Singes for all non-human primates.

Noteworthy, in 1811 Humboldt & Bonpland (1811) cited the still unpublished St.-Hilaire's (1812) descriptions of *M. humeralifer*, *M. melanurus*, *C. aurita*, *C. geoffroyi*, and *C. penicillata* but attributed to such author the authority of these names. Humboldt & Bonpland (1811) mentioned St.-Hilaire's (1812) classification of marmosets into genus *Jacchus* but considered all Platyrrhini in the genus *Simia* as Linnaeus (1758), Linnaeus (1758) created the family Hapales for marmosets, and included squirrel monkeys and titi monkeys in the family Sagoins.

Kuhl (1820), considering mainly teeth and ear characters, mixed the classifications of Illiger (1811), Humboldt & Bonpland (1811), and St.-Hilaire (1812): marmosets in the genus *Hapale* and family Hapales, squirrel and titi monkeys in the genus *Callithrix* and family Sagouin. Gray (1821) provided a family classification of marmosets and, for the first time, erected names with appropriate Latin declinations—although some misspellings. Such author considered the dental formula and tail prehensility to define the family Harpaladae (recte Hapalidae), containing the genus *Harpale* (recte *Hapale*) for *C. jacchus*, and the family Callitricidae (recte Callitrichidae), containing *Callitrix* (recte *Callithrix*) for *Saimiri* cf. *sciureus*, *Sapajus apella*, *Ateles paniscus*, and *A.* cf. *belzebuth*. Hapalidae is therefore the first valid family name for marmosets and tamarins as correctly pointed out by Hershkovitz (1977) and Groves (2008).

In 1823 Spix (1823) described the first pygmy marmoset species and adopted the genus *Jacchus* to classify this and all other marmosets on the basis of tail anatomy. This author used *Callithrix* for titi monkeys and included this genus along with *Jacchus* in the Trichuri family. Wied-Neuwied (1826) collected marmosets on the Atlantic Forest of southeast Bahia State, Brazil, which were considered as *C. penicillata* for more than 150 years until Coimbra-Filho (1986) described them as belonging to a separate species, *C. kuhlii*.

Two decades later Lesson (1840) described two new *Hapale* subgenera for marmosets based on ear hairiness and tail pelage: *Hapale* (*Mico*) for the species with exposed ears (*M. argentatus*, *M. melanurus*), and *Hapale* (*Hapale*) for species with hairy ears or with a mane concealing ears and a ringed-pattern of tail pelage (*C. jacchus*, *C. aurita*, *C. penicillata*, *C. geoffroyi*, *Cebuella pygmaea*, *M. humeralifer*). *Mico* is therefore the priority genus-level name for Amazonian marmosets (see also Rylands et al, 2000). Although Wagner (1840) cited *Liocephalus* as a subgenus of *Hapale* for *M. argentatus* and *M. melanurus* (and other three tamarins) in the summary of his 1840's review, such name was not used since then and therefore is *nomen oblitum*. Lesson (1840) introduced the novelty of dividing marmosets into two separate groups and considered all primates as belonging to Simiadae family. Apart of Lesson (1840) and Wagner (1840), the classification of marmosets into subgenera remained largely unused.

Noteworthy, the description of subgenus *Mico* is vague and misleading: "Milieu de la face nu et aplati; favoris sur les joues et petit barbe sous le menton; poils généralement longs et soyeux; oreilles grandes et nues; queue longue et mince" (Lesson, 1840). 'Middle of the face naked and flat', and 'sideburns and a high concentration of hairs under the chin' are observed in the majority of Neotropical primates; 'hairs in general long and silky', 'big naked ears', and 'long and slender tail' are all characteristics observed virtually in all tamarins and most of the marmosets.

Wagner (1842) subsequently described *Harpale chrysoleucos* (recte *Hapale*) [=*Mico chrysoleucos* (Natterer, 1842)] attributing such name's authority to Johann Natterer, to whom in fact the name belongs to. In 1865 Gray (1865) corrected the spelling of his genus *Harpale* and followed the rationale of Lesson (1840) (ear hairiness and color of tail pelage) to separate marmosets into four species groups: Hapale (for *C. aurita*), Jacchus (for *C. jacchus*), Cebuella (for *Ce. pygmaea*), and Mico for (*M. argentatus* and *M. melanurus*, considered this as a single species). Gray (1865) described the Mico species group as marmosets having "...ears naked, exposed, without any ear-pencil. Tail uniformly black".

Gray (1868) posteriorly described *Mico sericeous*, a junior synonym of *Mico chrysoleucos*, which has tufted ears and golden-white ringed tail—therefore contradicting his previous definition of species in Mico group. Gray (1870) formally elevated his four *Hapale* species groups to genus level and created a new genus, *Micoella*—in which he included *Micoella chrysoleucos*, another junior synonym of *Mico chrysoleucos*—adopted the genus *Callithrix* for titi monkeys, and classified all Platyrrhini in Cebidae family. According to Gray

(1870), marmosets were divided into five genera: *Hapale*, *Jacchus*, *Cebuella*, *Mico*, and *Micoella.*

The period spanning the initial investigations of morphological relationships of marmosets in the 17th century to the 19th century generated a profusion of different genus names, as well as a plethora of classifications schemes (see also Hershkovitz, 1977; Groves, 2008). The marmoset systematics shifted from using a single genus—that included also tamarins and saki monkeys—to up to five genera. The use of subgenera, as proposed by Lesson (1840) did not find acceptance by researchers and such subcategory is uncommon in the classification of marmosets. The use of the same genera for marmosets, squirrel and titi monkeys also was very common. The classification of marmosets (and tamarins) at the family level had started to incorporate correct spellings and declinations, although these were confusing in nomenclature and incongruent in composition.

In the beginning of the 20th century Thomas (1903a,b) started to clarify the usage of genus names for marmosets, adopting back a single genus—*Callithrix*—for Atlantic forest, Amazonian, and pygmy marmosets; this author also described *C. flaviceps*. Thomas (1903b) argued for the use of Callitrichidae [=Callitricidae Gray, 1821] for marmosets and tamarins, although Gray (1821) did not include any of these taxa in such family. Gray (1821) included *C. jacchus* – the only marmoset or tamarin considered in his review – in Harpaladae (*recte* Hapalidae) family. Thomas (1904) subsequently described the Goeldi's monkey as *Midas goeldii* and Miranda-Ribeiro (1912) described a junior synonym of this species, *Callimico snethlageri*. The genus *Callimico* from Miranda-Ribeiro (1912) has been used since then and *Cl. goeldii* is the only species of this genus known to date.

Nonetheless, the confusion on the usage of genus and family names and classifications for marmosets continued. Based on pelage patterns, geographical distributions, skull and teeth anatomy, Elliot (1913) classified Amazonian, Atlantic forest, and pygmy marmosets plus the

Goeldi's monkey into genus *Callithrix*, and included it in the family Callitrichidae along with titi monkeys and tamarins. Pocock (1917) relying on the anatomy of hands, feet, ears, and teeth, classified all marmosets in the genus *Hapale*, family Hapalidae. Pocock (1920, 1925) subsequently mentioned Goeldi's monkeys as *Cl. goeldii* in the Callimiconinae subfamily of Hapalidae. Elliot (1913) and Pocock (1917, 1920, 1925) provided the first comprehensive studies on comparative anatomy of marmosets and the Goeldi's monkey.

Thomas (1920) changed his previous classification (Thomas, 1903b) by adopting *Hapale* for marmosets, and described a new species—the Snethlage's marmoset *M. emiliae*. Thereafter Thomas (1922) provided a classification on the basis of pelage pigmentation of tail and limbs, and on the presence or absence of ear tuffs in three genera: *Hapale* for tufted-ear marmosets, *Mico* for untufted-ear marmosets, and *Cebuella* for pygmy marmosets. Thomas (1922) also described *M. leucippe* and classified the Goeldi's monkey in *Callimico*. In 1940, the second pygmy marmoset was discovered and described as *Cebuella niveiventris* by Lönnberg (1940). Both northern and southern pygmy marmosets, *Ce. pygmaea* and *Ce. niveiventris*, are currently recognized as full species—the only known species for this genus (Boubli et al, 2021).

Cruz-Lima (1944) used the genera *Cebuella* and *Callithrix* to classify marmosets based on teeth and jaw anatomy, both in Callitrichidae family, and classified Goeldi's monkey in as genus *Callimico*, subfamily Callimiconinae of Cebidae. Simpson (1945) adopted the genus *Callithrix* for all marmosets and grouped these with tamarins in Callitricidae (recte Callitrichidae) family while all other Platyrrhini were grouped in Cebidae––*Callimico* as subfamily Callimiconinae of the latter Hill (1957), based on a comparative anatomy study, adopted *Mico* for *M. argentatus* and *Cebuella* for pygmy marmosets, keeping the other marmosets in *Hapale*, therefore relying on three genera for the classification of marmosets as did Thomas (1922). Hill (1957) used Hapalidae family for marmosets and tamarins, and Callimiconidae for *Cl. goeldii*.

Based on a literature review, Cabrera (1957) classified pygmy marmosets in genus *Cebuella* and all other marmosets in *Callithrix*, both in Callitrichidae family—with Goeldi's monkey in genus *Callimico*, Callimiconinae subfamily of Cebidae. Subsequently, Cabrera & Yepes (1960) classified tufted-ear marmosets in *Hapale*, untufted-ear marmosets in *Mico*, and pygmy marmosets in *Cebuella*, all in the subfamily Hapalinae of Hapalidae. According to these authors, marmosets should be classified in three genera, as proposed by Thomas (1922), and classified *Cl. goeldii* in the subfamily Callimiconinae of Hapalidae.

In Napier & Napier's (1967) review, the authors adopted *Cebuella* and *Callithrix* to classify marmosets, *Callimico* for Goeldi's monkey, all these in the Callitrichidae family. Ávila Pires (1969) followed the scheme of Napier & Napier (1967) for marmosets after the examination of museum specimens but included *Callimico* in the subfamily Callimiconinae of Cebidae. At this point there was a clear trend of classifying marmosets into two genera according to morphological relationships and *Callimico* was often classified in its own family, Callimiconidae, or as a particular subfamily of Cebidae.

This trend culminated in Hershkovitz (1977) who, based on a literature review plus a new, multisource morphological dataset on marmosets, tamarins, and Goeldi's monkey adopted *Cebuella* and *Callithrix* to classify marmosets, grouped these with tamarins (genus *Saguinus*) in Callitrichidae family, and *Cl. goeldii* in Callimiconidae. Nonetheless, this author distinguished two species groups within *Callithrix*: "argentata" for Amazon marmosets and "jacchus" for Atlantic Forest marmosets. Hershkovitz (1977) relegated most marmoset taxa to subspecies and described *M. intermedius*. Vivo (1991) revised the systematics of marmosets based on phenotypical characters and followed Hershkovitz's (1977) scheme but without distinguishing species groups and elevating all taxa to full species.

The 20th century testified an increased interest on morphological relationships of marmosets, achieving its maximum expression with Hershkovitz (1977). His work deeply influenced the systematics of marmosets and Goeldi's monkey and still find some acceptance to date (e.g. Rosenberger, 2020). That is, although most authors historically classified marmosets in distinct genera and the divergence between Atlantic Forest and Amazon marmosets was recognized even by Hershkovitz (1977), the trend for the two subsequent decades was to lump Atlantic Forest and Amazon marmosets in *Callithrix*, to classify pygmy marmosets in *Cebuella*, and *Cl. goeldii* in its own family or subfamily. However, this scenario started to change with the discovery of the dwarf marmoset and due to a new body of evidence on the evolutionary relationships of these primates from DNA-based phylogenetic inferences.

In 1998 Roosmalen et al. (1998) described the dwarf marmosets as *Callithrix humilis*, which was subsequently transferred to the genus *Mico* (Rylands et al, 2000) and then to a new genus, *Callibella* Roosmalen & Roosmalen, 2003 to avoid paraphyletic genera as the initial DNA phylogenies were showing Amazon marmosets as sister taxa to pygmy, and not Atlantic Forest marmosets (Schneider et al, 1993, 1996). Moreover, Aguiar & Lacher-Jr. (2003, 2009) and Marroig & Cheverud (2009) endorsed the distinction of dwarf marmosets in the genus *Callibella* based on skull and jaw morphology, and shown the disparity between Atlantic Forest, Amazonian, dwarf, and pygmy marmosets. Ford & Davis (2009) reinforced the distinction between these four marmoset groups based on post-crania characters and suggested a close relationship between Amazon marmosets and pygmy marmosets.

In the beginning of the 21st century the studies of comparative morphology incorporated a statistical background and achieved an increased accuracy and detail. It was also starting to become clear that pygmy, Amazon, Atlantic Forest, and dwarf marmosets are morphologically distinct. Moreover, it was nearly a consensus from morphological analyses that Goeldi's monkey was the first offshoot among marmosets and tamarins. At this point there were some phylogenetic studies based on morphological data and the phylogenetic studies based on molecular data were starting to flourish.

*The four decades of phylogenetic studies on marmosets and Goeldi's monkey*

Most hypotheses on the phylogenetic relationships of marmosets and the Goeldi's monkey available to date were proposed as result of broad studies targeting the evolutionary history of platyrrhines. Given the macro-scale of the analyses, these studies have low representativeness of specimens, species, and genera. Therefore, they are less informative than previous comparative morphology analyses focused on marmosets, Goeldi's monkey, or callitrichids—although most of those, in turn, did not adopt an explicit phylogenetic approach or cladistic analyses.

The first phylogenetic study including marmosets and Goeldi's monkey was based on immunological data: Cronin & Sarich (1975) revealed *C. jacchus* and *Ce. pygmaea* as sister taxa, the only marmosets included in the analyses, and Goeldi's monkeys as sister to them (see also Baba et al, 1979). Only two decades later the use of molecular data became more consistent in phylogenetic studies of platyrrhines. The seminal paper by Schneider et al. (1993) using sequences of the e-globin gene is the hallmark of an entire new generation of phylogenetic studies on Neotropical primates, based on DNA data. These authors showed *C. jacchus* and *Ce. pygmaea* as sister taxa, and *Cl. goeldii* as basal to them—lacking a representation of Amazon marmoset species. The relationships in Schneider et al. (1993) are poorly supported but the same topology was recovered with higher support in subsequent molecular phylogenies, e.g. Harada et al. (1995) using a larger dataset of e-globin gene sequences, Horovitz & Meyer (1995) using the 16S mitochondrial gene, and Schneider et al. (1996) using sequences of the e-globin and the IRBP intron 1 combined.

The first phylogenies based on morphology data relied on similar and partially shared datasets of teeth and skull characters. Such morphology-based trees generated hypotheses that (i) presumably all marmosets belong to a single lineage, classified into the *Callithrix* (Rosenberger, 1980; Rosenberger 1981) or that (ii) pygmy marmosets and presumably all other marmosets belong to two separate lineages, classified in to genera *Callithrix* and *Cebuella*, respectively (Rosenberger, 1984, 2020; Kay, 1990). Natori (1986) shown Atlantic Forest and Amazon marmosets as sister but separate clades according to dental characters, and these two as sister to the pygmy marmosets clade. It was nearly a consensus that *Callimico* was the first diverging lineage of Callitrichidae or closely related to tamarins and lion tamarins according to morphology-based phylogenies comparative morphological analyses (Hershkovitz, 1977; Rosenberger, 2020; Ford & Davis, 1992).

Whether based on morphology or molecular data the first phylogenetic studies on marmosets showed *C. jacchus* and *Ce. pygmaea* as sister species because the representativeness of marmoset species was very low and simply lacked Amazon marmoset species (*Mico*). Moreover, information on which specimens and species were analysed is absent in many of the first morphology phylogenies which, in many cases, impedes understanding whether *Callithrix* was used only for Atlantic Forest, for these and Amazonian marmosets, or for all marmosets. The Goeldi's monkey was retrieved as closely related to marmosets in molecular phylogenies, and as a first offshoot among Callitrichidae or as closely related to tamarins and lion tamarins in morphology-based phylogenies.

The first integrative phylogenetic analysis, based on molecular and morphology data then available, supported the sister taxon relationship of *Callithrix* and *Cebuella*, and Goeldi's monkey as a first Callitrichidae offshoot (Purvis, 1995). However, the analysis of Purvis (1995) was obviously biased towards morphology data because it shown Amazon and Atlantic Forest marmosets as sister groups, both in the genus *Callithrix*, pygmy marmosets as sister to these two, in genus *Cebuella*, and *Callimico* as the first Callitrichidae offshoot—all relationships with high support. Horovitz et al. (1998), using a combination of mitochondrial, nuclear, and morphological data also supported *C. jacchus* and *Ce. pygmaea* as sister taxa due to the lack of Amazon marmosets in the analysis, and shown Goeldi's monkey was sister to marmosets according to molecular data and as sister to tamarins according with morphology data.

Tagliaro et al. (1997) carried out the first phylogenetic inference focused for marmosets. Although analysing only a few species, individuals, and a single mitochondrial marker (dloop), these authors provided strong support for *Cebuella* as sister to *Mico* and not to *Callithrix*. Such relationships were recovered subsequently by Porter et al. (1997) and Goodman et al. (1998) both based on e-globin gene sequences, Nagamachi et al. (1999) based on cytogenetic data, Canavez et al. (1999a,b) based on ß2-microglobulin gene, Chaves et al. (1999) based on the intron 11 of the von Willebrand factor gene, Roosmalen et al. (2000) based on mitochondrial control region and intron 2 of the ß2-microglobulin gene, Tagliaro et al. (2000) based on mitochondrial gene ND-1, Chaterjee et al. (2009) based on seven mitochondrial genes (12s, 16s, COII, cytb, NADH3, NADH4L, and NDH4) and three nuclear genes (CXCR4, SRY, TSPY), and Canavez et al. (1996) based on chromosome morphology.

Barroso et al. (1997), however, inferred a phylogeny using the IRBP intron-1 that showed *Mico*, *Callithrix* and *Cebuella* in a trichotomy, and *Cl. goeldii* as sister to lion tamarins (*Leontopithecus rosalia*). Schneider (2000) using a combined dataset of four nuclear loci (ß2-microglobulin, IRBP, G6DP, and EPSILON) grouped Atlantic Forest and Amazon marmosets into *Callithrix*, retrieved as a sister taxon to *Cebuella*—Goeldi's monkey was retrieved as sister to the basal marmoset clade. Pastorini et al. (1998) specifically addressed the phylogenetic relationships of *Cl. goeldii* for the first time, using three tRNA markers and the ND4 mitochondrial gene. These authors showed *Cl. goeldii* as sister to marmosets but with conflicting levels of support depending on the method of phylogenetic inference and schemes used for data concatenation (Pastorini et al, 1998). Dornum & Ruvolo (1999) based on the G6PD locus, Opazo et al. (2006) using e-globin, IRBP, von Willenbrand factor, ß2-microglobulin, ß-globin, and G6PD loci, and Osterholz et al. (2009) based on *Alu* elements clearly showed the retroposon integrations and sister taxa relationship between *Cl. goeldii* and marmosets.

Horovitz (1999) further reinforced this relationship between *Cl. goeldii* and marmosets using the same molecular markers as Horovitz et al. (1998), but adding morphological data from fossils. This author showed that, in morphological grounds, Goeldi's monkey is closely related to extinct species—*Patasola*, *Carlocebus*, or *Branisella*—and that these, in turn, form a clade sister to marmosets according to molecular data. The close relationship between *Callimico* and marmosets was subsequently shown by Porter et al. (1997) but with low support using e-globin gene, Neusser et al. (2001) using cytogenetic data, Wildman et al. (2009) using a dataset of 28 loci, and Wang et al. (2018) using 56 nuclear non-coding loci. On the other hand, Garbino (2015), as in several previous morphology-based analyses, showed Goeldi's monkey as the first offshoot among Callitrichidae, or close related to tamarins/lion tamarins depending on the type of dataset considered in the analysis. Such view remains supported by some to date (Rosenberger, 2020).

The long-standing hypothesis that Atlantic Forest and Amazon marmosets should be classified into a single genus—*Callithrix*—and the phylogenetic position of *Callimico* as a first offshoot among callitrichids started to be challenged in the end of the 90's by molecular phylogenies including Amazon marmosets. Subsequently, the body of evidences for sister taxa relationship of *Mico* and *Cebuella*, with *Callithrix* as sister to both, and the Goeldi's monkey as sister to marmosets just increased. These relationships were recovered by Perelman et al. (2011), using a dataset of 54 nuclear genes from approximately 90% of the primate species, Aristide et al. (2015), who used 31 DNA loci, and Hugot (1998), who used molecular, morphological, and parasitological data. Rylands et al. (2009, 2012)—who had previously resurrected the genus *Mico* (Rylands et al, 2000) for Amazon marmosets to avoid paraphyly between *Callithrix* (Amazon plus Atlantic Forest marmosets) and *Cebuella*, based on the studies of Schneider et al. (1993, 1996)—reinforced the adoption of *Callithrix*, *Cebuella*, and *Mico* considering these additional evidences.

Nonetheless, the discovery of the dwarf marmoset by Roosmalen et al. (1998) brought further uncertainties on the relationships between marmoset lineages and their genus-level classification. Roosmalen & Roosmalen (2003) presented for the first time a hypothesis for the phylogenetic position of the dwarf marmoset and shown *Mico* as sister taxa to *Cebuella*, *Callibella* sister to both, all three forming a clade of low support. These authors also just proposed a new genus—*Callibella* Roosmalen & Roosmalen, 2003—for the classification of the dwarf marmoset.to form Cortés-Ortiz (2009) showed the same topology as Roosmalen & Roosmalen (2003) for *Mico*, *Cebuella*, and *Callibella*, and reinforced the sister taxa relationship of *Callimico* and marmosets. Roos & Zinner (2017) also shown *Mico* as sister to *Cebuella*, *Callibella* as sister to both, *Callithrix* as the first diverging lineage of marmosets, and *Callimico* as sister to the basal marmoset clade.

On the other hand, Schneider et al. (2012) using dloop sequences and four nuclear loci (*Alu* elements) shown *Mico* as sister taxa to *Callibella*, and *Cebuella* as sister to both with high support according to the dloop mitochondrion locus but with low support in the phylogeny of nuclear loci. The same topology was retrieved by Schneider & Sampaio (2015) using different sets of previously available DNA data, by Garbino (2015) using morphological data (but showing *Callimico* as sister to tamarins) and by Buckner et al. (2015) using four mitochondrial and six nuclear loci.

Silva et al. (2018) addressed for the first time the phylogenetic position of *Callibella* using genomic data. These authors showed convincingly that *Callibella* is sister to *Mico*, both sister to *Cebuella*. Boubli et al. (2018) also found this same topology using genomic data when inferring the relationships of *Cebuella* species. Nonetheless, none of these phylogenomic inferences considered a robust sampling in terms of specimens and species of all marmoset genera.

The 40 years of phylogenetic studies on marmosets and Goeldi's monkey can be summarized as a succession of hypotheses that are largely incongruent—even with the observed increase in the amount and quality of molecular data, and in the computational capacity to perform such analyses. These incongruencies are evident regardless the type of data used if one compares studies based on morphology only, based on molecular characters only, or studies based on both and other types of data. To this date, there was no clue on the reason for such incongruencies. Consequently, the genus and family classifications of marmosets and Goeldi's monkey so far have been more or less adopted according to personal or operational preferences rather than biological reasoning.

*An up-to-date systematic phylogenetic of marmosets and Goeldi's monkey*

In this paper we decisively recovered the evolutionary relationships of marmosets (*Callithrix*(*Cebuella*(*Mico*(*Callibella*)))) and demonstrate that incomplete lineage sorting is the main reason underlying incongruencies in previous studies, associated with the shallow taxonomic, individual and DNA sampling. We also show that Goeldi's monkey is sister to marmosets and point out that the main reason for the previous incongruencies in their classification and phyletic position is the previous use of homoplastic characters. Therefore, we provide a robust and stable phylogenetic background for the classification of marmosets at the genus level and for the Goeldi's monkey at family level.

Moreover, it is plenty of data on the biological disparities of marmosets—although the species of *Mico* and especially *Ca. humilis* are underrepresented in previous studies. These data concern variation in shape and size of skull, jaw, and postcrania, body size and mass, pelage and tegument, karyology, and zoogeography of the four marmoset lineages—the Amazon (*Mico*), the dwarf (*Callibella*), the pigmy (*Cebuella*), and the Atlantic Forest marmosets (*Callithrix*).

Aguiar & Lacher (2003) using 32 skull and jaw measurements showed a clear distinction between *Callibella*, *Cebuella*, and *Mico* in size and shape, related to different levels of gum-feeding in discriminant function analysis (DFA). Aguiar & Lacher (2009) using the same approach further reinforced the existence of four morphogroups that correspond to *Callibella*, *Callithrix*, *Cebuella*, and *Mico* genera, which are related to size and allometry in DFA; in the same study but considering a subset of 10 skull measurements the authors shown four morphogroups of marmosets corresponding to the four genera according to size. Marroig & Cheverud (2009) using 39 craniofacial characters derived from 3-d morphometrics and principal component (PCA), variance, and multivariate allometric size-scaling analyses found significant variation in size and allometry and showed three well-defined morphogroups corresponding to *Cebuella*, *Callithrix*, and *Mico*. Silva et al. (2018) using 11 skull measurements clearly shown that *Callibella*, *Mico*, and *Cebuella* form three distinct morphogroups related to size variation in PCA. Ford & Davis (2009) based on the analysis of 160 measurements of marmosets postcrania and DFA and ANOVA analyses found that *Callibella*, *Callithrix*, *Cebuella*, *and Mico* form four clearly distinct morphogroups, especially regarding shape and size of humerus, radius, ulna, and femur shape. Garbino (2015) also point out discrete variation in skull, postcrania, pelage and tegument of *Callibella*, *Callithrix*, *Cebuella*, and *Mico* although did not recognize *Callibella* as a valid genus.

There is a clear size-gradient from the smallest (*Cebuella*) to the largest marmosets (*Mico*). *Cebuella pygmaea* and *Ce. niveiventris* are the smallest monkeys worldwide (Hershkovitz, 1977), *Callibella humilis* is intermediary in body size and mass (Silva et al, 2018) between *Cebuella* and *Callithrix* (Hershkovitz, 1977), and *Mico* (RCA unpub. data) are larger than *Callithrix*. According to Hershkovitz (1977), Groves (2001), Roosmalen et al. (1998), Roosmalen & Roosmalen (2003), Garbino (2015), and Costa-Araújo et al. (2019, 2021) the pelage patterns differ consistently between the four marmoset lineages. And according to Nagamachi et al. (1992, 1994, 1997, 1999), Canavez et al. (1996), Oliveira et al. (2012), and Stanyon et al. (2018) the karyotype of *Callithrix*, *Cebuella*, and *Mico* are not only different but also highly conserved—most chromosome rearrangements are autapomorphic evolutionary events and have little levels of convergence. The karyotype of *Cb. humilis* was not studied to date.

Except for *Callibella*, each marmoset lineage occurs on a particular area within South America (see Figure 1). *Callithrix* species range along the Atlantic forest, Caatinga and Cerrado biomes east of South America, are endemic to Brazil and isolated from *Mico* species eastwards by the dry diagonal composed by the Caatinga and Cerrado biomes (Rylands et al, 2009; Rylands & Mittermeier, 2013; Culot et al, 2019). The *Mico* species are restricted to east of Madeira River, west of Tocantins River, and south of the Amazonas River in southeast Amazonia, with exception of *M. melanurus* which occurs also in a small part of Cerrado and Pantanal in Brazil (west to the range of *Callithrix penicillata*), and in Chaco in Bolivia and Paraguay (Rylands et al, 2009; Rylands & Mittermeier, 2013; Culot et al, 2019; Costa-Araújo et al, 2019, 2021). The range of *Cb. humilis* encompasses a small area on the Aripuanã-Machado interfluve, southcentral Amazonia, over the distribution of *Mico* (Silva et al, 2018), reaching south as far as the Igarapé Preto, Tenharin Indigenous land (Costa-Araújo, 2020). *Cebuella* species range northwest Amazonia and are limited by the Madeira River in the south and by the Japurá River in the north; the southernmost records are in Ponton, Bolivia, and in the Manu National Park, Peru; the westernmost record is Copataza River, Equator, and the northernmost record is Puerto Limón, Putumayo, Colombia (Boubli et al, 2018).

Taking all evidences together, after accounting for monophyly and clade stability—as demonstrated by our phylogenomic analyses—plus phenotypic diagnosability, zoogeography, nomenclature and nomenclatural stability, community consensus—as here revised—and the fact that these primates are among the more widely-known monkeys of the Neotropics (taxon naming criteria 1, 2, 3, 5, 9, 10, and 11; Vences et al, 2013), the adequate option for classification of the four marmoset clades is the four genera scheme: *Callibella*, *Callithrix*, *Cebuella*, and *Mico.* Each of these genera is monophyletic and the species therein are morphologically, anatomically, ecologically, chromosomally, ontogenetically, phylogenetically, and zoogeographically distinct. Therefore, such scheme fulfils the basic principle of any classification—which is to provide a universal, stable, and easy-to-adopt system of meaningful names and categories for research and communication on the natural entities such names and categories belong to. Moreover, this is a powerful and accurate system of information retrieval, as suggested by Groves (2004) a classification should be.

The adoption of *Mico* for Amazon marmosets, *Cebuella* for pygmy marmosets, and *Callithrix* for Atlantic Forest marmosets has indeed been widely accepted since this scheme was proposed by Rylands et al. (2000). Moreover, the IUCN's RedList (Mittermeier & Röhe, 2021) and primate specialist group (IUCN, 2021), and the majority of the research community (e.g. Roosmalen & Roosmalen, 2003; Aguiar & Lacher, 2003, 2009; Cortés-Ortiz, 2009; Ford & Davis, 2009; Rylands &Mittermeier, 2009, 2013; Rylands et al, 2009, 2012; Paglia et al, 2012; Roos & Zinner, 2017; Boubli et al, 2018, Silva et al, 2018; Costa-Araújo et al, 2019, 2021) has also considered *Callibella* as a valid genus. The disagreement on our proposal is restricted and based on two flawed arguments.

One of the arguments for not recognizing *Callibella* is based on divergence time, following the rationale of primate classification according to divergence time thresholds of Goodman et al. (1998). Schneider et al. (2012) and Schneider & Sampaio (2015) argued for the classification of *Ca. humilis* in genus *Mico* considering that their divergence time is of the same magnitude of the divergence between *C. aurita* and other *Callithrix* species. Nonetheless, divergence time or the thresholds proposed by Goodman et al. (1998) should not be used as a rule thumb rule criterion in primate taxonomy and systematics.

First, this approach has two conceptual errors: (i) it assumes that evolution operated at the same extent and depth across the primate tree of life, and (ii) considers a proxy of trait diversification rather than the diversification of the trait itself. Second, this approach carries an operational error that follows up the first conceptional error: it is clear that divergence times and the thresholds proposed by Goodman et al. (1998) to classify primate genera (4-6My) do not necessarily reflect morphological or ecological diversification in platyrrhines. This fact is observable, on one hand, in the divergence times of *Mico* and *Callibella* (1.75My) and their distinctions as described above, and of *Cebus* and *Sapajus* (2.11My) and their well-known distinctions (Lynch-Alfaro et al, 2012); and, on the other hand, on the divergence times and long-known similarities between *C. aurita* and the other *Callithrix* species (1.62My), and the two *Cebuella* species (1.45My; Hershkovitz, 1977; Soini, 1993) .

The other claim is based on a phylogenetic inference that did not offer confidence on the phylogenetic position of *Callibella* as sister clade to *Mico* species, due to low bootstrap support (Garbino, 2015). However, such relationship has been repeatedly recovered with maximum support with the use of genetic (Schneider et al, 2012; Schneider & Sampaio, 2015) and genomic data (Silva et al, 2018; Boubli et al, 2018; Costa-Araújo et al, 2019, 2021, this paper). More recently, after accepting the well-established phylogenetic position of *Callibella*, some authors further claimed that there were no other evidences for recognizing it as a valid genus and suggested its classification as a subgenus of *Mico* (Garbino et al, 2019). However, as summarised here, (i) there are divergences in the shape and size of skull, jaw, and postcrania, as well as ecological divergences between *Mico* and *Callibella*, and (ii) the classification of one (or some) marmoset lineage as subgenus is not phylogenetically coherent, is highly arbitrary, and bring unnecessary nomenclatural complexity to marmoset systematics. Therefore, the claims for not recognizing *Callibella* as a full genus based on divergence times and supposed lack of dissimilarities are flawed. Therefore, in addition to proposing the four genera classification scheme for marmosets, we below redescribe the genus *Mico* and provide updated definitions of *Callibella*, *Callithrix*, and *Cebuella* in order to pave the way for marmoset systematics usability.

We found that the incongruencies in the classification and phylogenetic position of *Callimico* are basically due to homoplasy of the characters traditionally used in past studies. Most of molecular-based phylogenies recovered *Cl. goeldii* as sister to marmosets, which is the same relationship we obtained here with genomic data for the entire Callitrichidae family. According to morphological data from fossils and subfossils, *Callimico* is the only living representative species of a more diverse clade composed by taxa now extinct (Horovitz, 1999). Therefore, there is no reason from an evolutionary perspective for the classification of *Callimico* in its own family—nor in a particular subfamily of Cebidae—unless other families or subfamilies, respectively, are raised for marmosets, tamarins, lion tamarins, capuchins, and squirrel monkeys. Such scheme, in our opinion, also bring only more unnecessary and useless complexity to the systematics and nomenclature of Callitrichidae.

*The redefinition of Mico and updated definitions of Callibella, Callithrix, and Cebuella*

Order Primates Linnaeus, 1758

Family Callitrichidae Gray, 1821

Redescription.

**Genus *Mico* (Lesson, 1840). Amazon marmosets.**

*Hapale* (*Mico*) Lesson, 1840: 192. "Mico" Buffon & Daubenton, 1756: 121; *Simia* Linnaeus, 1771: 521, part; Humboldt & Bonpland (1811): 359, part. *Callithrix* Erxleben, 1777: 55, part; Thomas (1903b): 457, part; Elliot (1913): 217, part; Simpson (1945): 65, part; Cabrera (1957): 185, part; Cruz-Lima (1957): 243; Napier & Napier (1967): 346, part; Ávila-Pires (1969): 50, part; Hershkovitz (1977): 480, part; Rosenberger (2020): 37, part; Schneider et al. (1993): 235, part; Schneider & Rosenberger (1996): 6, part; Vivo (1991): 21, part. *Sagoinus* Kerr, 1792: 80, part. *Hapale* Illiger, 1811: Kuhl (1820): 46, part; Wagner (1847): 463, part. Gray (1870): 63, part; Pocock (1917): 257, part; Thomas (1920): 269; Thomas (1922): 198, part; Pocock (1925): 38, part; Hill (1957): 280, part. *Hapale* (*Hapale*) Lesson, 1840: 185, part. *Jacchus* St.-Hilaire, 1812: 120, part. *Midas* St.-Hilaire, 1828-10: 36, part. *Liocephalus* Wagner, 1840: v, part. *Micoella* Gray, 1870: 130.

*Type species. Mico argentatus* (Linnaeus, 1771) by monotypy.

*Definition.* Lower jaw canine and incisive roughly of the same length, four pairs of molars, claw-like nails on all digits except the hallux, non-prehensile, slender tail, mostly restricted to the southern portion of Amazonia (south of Amazonas and east of Madeira Rivers) in Brazil but *M. melanurus* is also present on the northern Cerrado, Amazonia-Cerrado ecotone, Pantanal, and on the Bolivian and Paraguayan Chaco; the more species rich genus of marmosets, represented by a clade composed of 15 species currently recognized that is sister to the *Callibella* clade; the largest species among marmosets, ears nearly bare, with few, sparse, thin and short hairs in most of the species, ears lightly covered with long but few and sparse hairs (*M. intermedius*), or with ear tufts outgrowing from inner and outer pinnae (*M. chrysoleucos, M. humeralifer*, *M. mauesi*), mane absent, pale hip patches present (*M.

*acariensis*, *M. melanurus*, *M. humeralifer*, *M. mauesi*), saddle never striated, and only three species (*M. chrysoleucos*, *M. humeralifer*, *M. mauesi*) show a ring-like pattern in tail pelage due to bi-banding hairs, otherwise blackish, white, or yellowish in the other 12 species; malleolar orbicular apophisis present, genitals complex anatomically and hypertrophied in relation to body size, lingual cingulum of superior molar teeth continually extended mesial and distally around the protocone, metaconid of premolar 3, lateral process of pterigoid ending close to oval foramen, petrosal spine of the tympanic bullae well developed (Hershkovitz, 1977; Rylands et al, 2009; Garbino, 2015; Costa-Araújo 2020; Costa-Araújo et al, 2019, 2021).

*Species.* *Mico acariensis* (Roosmalen, Roosmalen, Mittermeier & Rylands, 2000), *Mico argentatus* (Linnaeus, 1771), *Mico chrysoleucos* (Natterer in Wagner,1842), *Mico emiliae* (Thomas, 1920), *Mico humeralifer* (St.-Hilaire in Humboldt & Bonpland, 1811), *Mico intermedius* (Hershkovitz, 1977), *Mico leucippe* Thomas, 1922, *Mico mauesi* (Mittermeier, Schwarz & Ayres, 1992), *Mico marcai* (Alperin, 1993), *Mico melanurus* (St.-Hilaire in Humboldt & Bonpland, 1811), *Mico munduruku* Costa-Araújo, Farias & Hrbek, 2019, *Mico nigriceps* (Ferrari & Lopes, 1992), *Mico rondoni* Ferrari, Sena, Schneider & Silva-Jr., 2010, *Mico saterei* (Silva-Jr. & Noronha, 1998), *Mico schneideri* Costa-Araújo, Silva-Jr., Boubli, Rossi, Hrbek & Farias, 2021.

*Body size and mass.* The largest species among the four marmoset genera; mass $\bar{x}$=372g; head and body: $\bar{x}$=233cm; tail: $\bar{x}$=329mm (Costa-Araújo unpub. data).

*Skull and jaw.* Compared to the other three marmoset genera, *Mico* species have straighter, less acute jaw base, with the lobe of the angular process extending only minimally below the gnathion, the mandible is strong, the horizontal ramus is straight, less arcuate (Aguiar & Lacher 2003, 2009), and the zygomatic orbital margin is large (Natori, 1994).

*Postcrania.* In comparison to the other three marmoset genera, *Mico* species have significantly narrowed medio-lateral dimensions of the distal humerus (including a narrow capitulum, narrow trochlear groove or gutter, and narrow medial epicondyle), and a narrow and short radial neck, indicating a highly narrowed and compressed elbow joint, that may facilitate increased vertical orientations of arm movements towards an increased exudate-feeding (Ford & Davis, 2009).

*Pelage and tegument.* The saddle is never striated and only three species (*M. chrysoleucos*, *M. humeralifer*, *M. mauesi*) show a ring-like pattern in tail pelage due to bi-banding pattern of hairs instead of true tail rings; the tail in the other 12 *Mico* species are blackish, white or yellowish; pale hip patches are exclusive of this genus and present in three species (*M. melanurus*, *M. humeralifer*, *M. mauesi*); ear tuffs are present in three species (*M. chrysoleucos*, *M. humeralifer*, *M. mauesi*) but outgrow always from both outer and inner pinnae, which are otherwise nearly bare in the other 11 species, mane absent, no blaze and the muzzle are naked (Hershkovitz, 1977; Groves, 2001; Roosmalen et al, 1998; Roosmalen & Roosmalen, 2003; Garbino, 2015; Costa-Araújo, 2020; Costa-Araújo et al, 2019, 2021).

*Karyotype.* Diploid number is 44, being 10 acrocentric, 32 metacentric or submetacentric, a submetacentric X and an acrocentric or metacentric Y, both 1/10 and 2/16 syntenic associations with human chromosomes are present, high amount of heterochromatin distributed all over chromosomes' regions; relatively to *Callithrix*, a Robertsonian-type fusion of chromosomes 20 and 16, and a paracentric inversion in chromosome 19 (Barros et al, 1990; Nagamachi et al, 1992, 1994, 1999; Canavez et al, 1996; Stanyon et al, 2018).

*Geographical distribution.* East of Madeira and Guaporé Rivers, west of lower Tocantins and Araguaia Rivers, and south of Amazonas River, across the States of Amazonas, Pará, Mato Grosso, and Rondônia in the Brazilian Amazonia, extending to the northern part of the Brazilian Cerrado and Pantanal, and a small stretch of the northern Chaco in Bolivia and

Paraguay, west to the range of *Callithrix penicillata* (Rylands et al, 2009; Rylands & Mittermeier, 2013; Culot et al, 2019; Costa-Araújo, 2020).

*Evolutionary history.* The *Mico* lineage arouse during the Quaternary, around 1.75 Mya with the split of the *Mico*+*Callibella* ancestor on southcentral Amazonia and underwent a speciose and fast radiation of 15 species known to date in the past 700ky; these species are nested in four intra-generic clades or lineages here named for the sake of convenience: "red-face marmosets", wherein the six species usually present a reddish tegument on face (*M. argentatus*, *M. emiliae*, *M. leucippe*, *M. intermedius*, *M. munduruku*, *M. rondoni*); the "black-face marmosets", wherein all species present variable levels of eumelanism of the face tegument (*M. acariensis*, *M. marcai*, *M. melanurus*, *M. nigriceps*, *M. schneideri*); the "hairy-ear marmosets", wherein the three species possess ear tufts outgrowing from inner and outer pinnae (*M. chrysoleucos*, *M. humeralifer*, *M. mauesi*); and the "Sateré–Mawé marmoset", a single-species clade formed by *M. saterei*. The hairy-eared marmosets' clade was the first to branch off around 630kya, followed by the branching off of the Sateré–Mawé clade; the red-faced marmosets' clade is sister to the black-faced marmosets' clade and both split off 200kya.

*Ecology.* Inhabit *Terra Firme*, *Campinarana* (white sand savanna patches), *Vargem* (seasonally flooded forests along rivers), and *Várzea* (in permanently flooded forests) ecosystems of Amazonia, the Amazonia–Cerrado ecotone, and forest ecosystems of Cerrado, Pantanal, and Chaco, on old-growth, second-growth, and logged forests, forest edges and clearings, patches of exotic tree monoculture (such as *Tectona grandis*), and in forests fragments surrounded by rural and urban landscapes. *Mico* and *Callibella* taxa are sympatric, which is the sole case of syntopy without hybridization in marmosets (Rylands & Mittermeier, 2013; Costa-Araújo, 2020).

*Etymology and key applications of Mico.* According to Gumilla (1741), “mico” is a noun created by the “natives” of the Orinoco basin, Colombia, to designate the smallest monkeys on that region—therefore, it is originally a reference to a tamarin as none *Mico* species occurs on this region. Buffon & Daubenton (1756) transformed such tamarin’ popular name into a marmoset’ vernacular name when described *Mico argentatus* as “Le Mico”, before the publication and establishment of the Linnean system. Lesson (1840) transformed such popular name into a scientific name by creating the subgenus *Hapale* (*Mico*) for *M. argentatus* (and also *M. melanurus*, considered as an ‘old variant’ of the first). Gray (1865) adopted *Mico* to name a species group for Amazon marmosets, to describe *M. sericeous* (Gray, 1868), and then as a genus name (for *M. melanurus*) within a classification scheme (Gray, 1870). *Mico* was relatively often accepted until Hershkovitz (1977) decree *Callithrix* for all marmosets, then resurrected in the 2000’s as a genus name for the Amazon marmosets by Rylands et al. (2000), which have been widely accepted since then.

*Vernacular names.* saguis-da-Amazônia (Português), titís de Amazonia (Español), Amazon marmosets (English), Amazon Seidenäffchen (Deutsch), Ouistitis Amazoniens (Français).

**Genus *Callibella* Roosmalen & Roosmalen, 2003. Dwarf marmosets.**

*Callibella* Roosmalen & Roosmalen, 2003: 2. *Callithrix* Erxleben, 1777: Roosmalen et al. (1998): 8; Rosenberger (2020): 37, part. *Mico* (Lesson, 1840): Schneider et al. (2012): 6, Schneider & Sampaio (2015): 7, Garbino (2015): 661. *Mico* (*Callibella*): Garbino et al. (2019): 2.

*Type species.* *Callibella humilis* (Roosmalen, Roosmalen, Mittermeier & Fonseca, 1998) by original designation.

*Definition.* Very small-bodied marmosets (mass $\bar{x}$=131g; head–body $\bar{x}$=15cm; tail $\bar{x}$=23cm), slightly larger than *Cebuella* species, non-prehensile, slender tail, ears exposed with white hairs outgrowing from the centre of pinnae, white rim around the face, mane absent, restricted to the southern portion of the Brazilian Amazonia (south of Amazonas and east of Madeira Rivers), in sympatry with *Mico*; canine and incisive of the same length on the lower jaw; four pairs of molars; muscular line delimits the middle third of the surface of the parietal bone, in a caudal–rostral direction, continuous and rougher in the frontal bones; the temporalis muscle is small and covers approximately 2/3 of the parietal bone and a small portion of the lateral surface of the frontal bone; the external occipital protuberance in the middle third of the occipital bone is longilineus laterally (Roosmalen et al, 1998; Roosmalen & Roosmalen, 2003; Garbino, 2015; Silva et al, 2018; Costa-Araújo, 2020).

*Species.* *Callibella humilis* (Roosmalen, Roosmalen, Mittermeier & Fonseca, 1998).

*Body size and mass.* Intermediary between *Cebuella* and *Callithrix* (mass: $\bar{x}$=131g; head and body: $\bar{x}$=156cm; tail: $\bar{x}$=237mm; Silva et al, 2018)

*Skull and jaw.* Compared to the other marmosets the jaw is less gracile, condyle is low, the condylion lies marginally above the occlusal plane, intermediate coronoid, the coronion is not as bold and high as in *Callithrix* but more developed than the sharp light hook of *Cebuella*—combined with a uniquely protruding angular process projected even broader and deeper than in *Cebuella* (Aguiar & Lacher, 2003, 2009).

*Postcrania.* Small femoral head, narrow femoral lateral condyle and tibial plateau, a short astragalar body with a short posterior facet for the calcaneus, short calcaneus with a short anterior extension, very narrow posterior trochlear width on the humerus, a tall anterior rim on the proximal radius for the ulna, and a wide calcaneal posterior facet, suggesting a less specialized clinging but marked similarities to *Cebuella*, which may represent a convergent evolution of those, associated with size reduction (Ford & Davis, 2009).

*Pelage and tegument.* Pelage of adults is entirely brownish with exception of the blackish tail; the mane and ear tuffs are absent but white hairs grow from the centre of pinnae, the crown is black and triangular, white rim around a naked face (Roosmalen et al, 1998; Roosmalen & Roosmalen, 2003; Garbino, 2015; Costa-Araújo, 2020).

*Geographical distribution.* The range of *Cb. humilis* encompasses a small area on the Aripuanã-Machado interfluve, southcentral Amazonia, over the distribution of *Mico*, reaching south as far as the Igarapé Preto, Tenharin Indigenous land, east of Madeira River and south of Amazonas River, south–central Amazonia, Brazil (Silva et al, 2018; Costa-Araújo, 2020).

*Evolutionary history.* The *Callibella* lineage arouse during the Quaternary, around 1.75 Mya, with the split of the *Mico+Callibella* ancestor on southcentral Amazonia as result of ecological speciation and character displacement.

*Ecology.* *Callibella* occur in sympatry with *Mico*, which is the sole case of syntopy without hybridization in marmosets, inhabiting *Terra Firme*, *Igapó*, and *Campinarana* ecosystems of Amazonia (Roosmalen & Roosmalen, 2003; Silva et al, 2018).

*Etymology.* "calli" derives from the Greek word kalos, and "bella" is an Italian word, both mean beautiful (Roosmalen & Roosmalen, 2003).

*Vernacular names.* saguis-anões (Português), Dwarf marmosets (English), tití enano (Español), Zwerg Seidenäffchen (Deutsch), Ouistiti nain (Français).

**Genus *Callithrix* Erxleben, 1777. Atlantic forest marmosets.**

*Callithrix* Erxleben, 1777: 55. *Callitrix* Boddaert, 1784: 42. *Sagoinus* Kerr, 1792: 80. *Sagouin* Lacepède, 1799: 4. *Saguin* Fischer, 1803: 113. *Sagoin* Desmarest, 1804: 8. *Hapale* Illiger, 1811: 71, part.; *Saguinus* Illiger, 1811: 71. *Jacchus* Humboldt & Bonpland, 1811: 359. *Arctopithecus* Cuvier, 1817: 115. *Harpale* Gray, 1821: 298. *Jacchus* Spix, 1823: 32, St.-Hilaire 1828-10: 36 in part *Ouistitis* Burnett, 1828: 307. *Hapales* Cuvier, 1829: 401.

*Antopithecus* Cuvier, 1829: 401. *Jachus* Schlegel, 1876: 254. *Sagoninus* Pocock, 1917: 257. *Cuistitis* Cabrera, 1958: 185.

*Type species.* *Callithrix jacchus* (Linnaeus, 1758) by subsequent designation (Thomas, 1903: 457).

*Definition.* Auricular tuffs inserted in front, around or on the inner surface of pinnae, saddle and rump with grey-black to brownish striated pelage, white blaze and hairy muzzle; lower jaw incisive and canine roughly of the same size; front-nasal profile convex; I2 smaller than I1; manubrium head of sternum in trapezoid shape; manubrium tail of sternum pronouncedly constricted, medial epicondile of humerus size of the medial epicondyle of the humerus, laterally projected, perpendicular to humerus axis; petrosal spine little developed, M2 reduced in comparison to M1, M1 entocingulum mesially and lingually interrupted entocingulus envolving distal protocone in M1, lingual cingulus underdeveloped in P4, high intermembral index, wide radial head, medio-laterally wide ulnar notch, tall cuboid facet on the calcaneous (Natori, 1986; Hershkovitz, 1975, 1977; Vivo, 1991; Garbino, 2015; Ford & Davis, 2009).

*Species.* *Callithrix aurita* (St.-Hilaire in Humboldt & Bonpland, 1811), *Callithrix flaviceps* (Thomas, 1903), *Callithrix geoffroyi* (St.-Hilaire in Humboldt & Bonpland, 1811), *Callithrix jacchus* (Linnaeus, 1758), *Callithrix kuhlii* Coimbra-Filho, 1986, *Callithrix penicillata* (St.-Hilaire in Humboldt & Bonpland, 1811).

*Body size and mass.* Smaller than *Mico* species; mass: $\bar{x}$=300g; head and body: $\bar{x}$=220cm; tail: $\bar{x}$=307mm (Hershkovitz, 1977).

*Skull and jaw.* Compared to species of the other three marmoset genera the lower jaw is sturdy, shows a less aggressive coronion, a condylar pivot which lies nearer to the occlusal plane, strongly recurved inferior margin of horizontal ramus, and deep angular lobe (Aguiar & Lacher 2003, 2009), small zygomatic orbital margin (Natori, 1994).

*Postcrania.* Compared to species of the other three marmoset genera, a wide radial head, a medio-laterally wide ulnar notch, and a tall cuboid facet on the calcaneus; widest medial epicondyles among callitrichids, the least specialized on vertical clinging among marmosets (Ford & Davis, 2009).

*Pelage and tegument.* The saddle is always transversely striated, a strongly ringed grey and black tails, the ear tuffs are always present and inserted in front (*C. penicillata* and *C. geoffroyi*), around (*C. jacchus*), or in a small area of inner pinna (*C. flaviceps* and *C. aurita*), whereas the mane—present in *Cebuella* species—and the whitish hip patches—present in *Mico* species—are absent, and the face has a white blaze and hairy muzzle (Hershkovitz, 1977; Groves, 2001; Garbino, 2015).

*Karyotype.* Diploid number is 46, being 14 acrocentric, 30 metacentric or submetacentric, a submetacentric X, and a metacentric, submetacentric or acrocentric Y, syntenic association with human chromosomes 1/10 present and the 2/16 absent, amount of heterochromatin is small and restricted to centromeres (Canavez et al, 1996; Nagamachi et al, 1997; Stanyon et al, 2018).

*Geographical distribution.* The six species range along the Atlantic Forest, Caatinga, and Cerrado biomes at east of South America, are endemic to Brazil, and mainly isolated from *Mico* species by the dry diagonal composed by the Caatinga and Cerrado biomes (Rylands et al, 2009; Rylands & Mittermeier, 2013; Culot et al, 2019; Malukiewicz et al, 2021).

*Evolutionary history.* Among marmosets the *Callithrix* lineage was the first to arouse, during the middle Pliocene (3.77Mya) in the Atlantic Forest with subsequent speciation events occurring in the Quaternary, from at least 1.62My (*C. aurita*) to 470Ky (*C. geoffroyi* and *C. kuhlii*).

*Ecology.* The species in general eat primarily exudates, and also invertebrates, fruits and other plant parts, small vertebrates, and fungi; group sizes vary from 2-15 individuals, home range sizes vary from 2 to 138.5ha, the groups can travel from 480 to 1980m per day, in a myriad of different ecosystems such as restinga, mangrove, Atlantic Forest evergreen, semi-decidual, decidual forests, Cerrado forests and gallery forests, Caatinga forests, secondary, mature, disturbed, logged and forest edges, orchards, urban areas and gardens, tree monocultures (Malukiewicz et al, 2021).

*Etymology.* "calli", derived from kalos, means beautiful and thrix means hair, both Greek words (Barnett, 2004).

*Vernacular names.* saguis-da-Mata-Atlântica (Português), titis del bosque Atlantico (Español), Atlantic Forest marmosets (English), Seidenäffchen (Deutsch), Ouistiti dans le Atlantic forêt (Française).

**Genus *Cebuella* Gray, 1865. Pygmy marmosets.**

*Cebuella* Gray, 1865: 734. *Callithrix* Erxleben, 1777: 55, part; *Hapale* Illiger, 1811: 71, part; Wagner (1847): 463, part; *Jacchus* Spix, 1823: 32, part; *Midas* St.-Hilaire, 1828: 36, part.

*Type species.* *Cebuella pygmaea* (Spix, 1823) by monotypy.

*Definition.* (mass $\bar{x}$=125g; head–body $\bar{x}$=13cm; tail $\bar{x}$=20cm)

Naked ears entirely concealed by mane, ear tuffs absent, elevated intermembral index, the saddle is brown black tabby, tails are all brown but can be slightly ringed agouti, and they bear a white moustache or dots by nostrils; condylar process at the same level or slightly below the teeth line, angular process ventrally projected into the horizontal ramus, protocone absent in P2, reduction index of the upper molar 2, pterygoid shape, height of mandibular condyle relative to planum alveolar, protocone of premolar 2 (Hershkovitz, 1977; Aguiar & Lacher, 2003; Garbino, 2015)

*Species.* *Cebuella pygmaea* (Spix, 1823), *Cebuella niveiventris* Lönnberg, 1940.

*Body size and mass.* The smallest monkeys worldwide. Mass: $\overline{x}$=125g; head and body: $\overline{x}$=136cm; tail: $\overline{x}$=202mm (Hershkovitz, 1977).

*Skull and jaw.* Lower jaw is gracile, condyle is low, condylion lies directly in line with the tops of the molar and premolar teeth, resting within their occlusal plane, sharply angled coronoid, deep angular lobe but very gracile (Aguiar & Lacher 2003, 2009). The zygomatic orbital margin in *Cebuella* is intermediate between *Callithrix* and *Mico* (Natori, 1994).

*Postcrania.* Many features in the femur, tibia, astragalus, and calcaneus are significantly reduced in size, including femoral head dimensions, several condylar and tibial plateau dimensions, width of the astragalar trochlea, several ventral astragalar dimensions, length of the calcaneus and of its anterior portion and posterior facet, length of the cuboid, of the fourth metatarsal, and of the combined power arm of the foot—in addition, the intermembral index is low, reflecting a shortened hindlimb/long forelimb; the most specialized on vertical clinging, especially in hindlimb morphology likely reflecting higher frequencies of scansorial and clinging behaviour and higher dependence on exudates (Ford & Davis, 2009).

*Pelage and tegument.* Mane is present, completely enclosing pinna, the saddle is brown black tabby, tails are all brown but can be slightly ringed agouti, and they bear a white moustache or dots on nostrils (Hershkovitz, 1977; Groves, 2001; Garbino, 2015).

*Karyotype.* Diploid number is 44, being 10 acrocentric, 32 metacentric or submetacentric, a submetacentric X and an acrocentric Y (Nagamachi et al, 1992), and almost identical to the karyotype of *Mico*. The main karyotypic distinction between such genera lies on the amount and distribution of heterochromatin, which is pronounced and widespread in chromosomes of *Mico* species (Nagamachi et al, 1992). The difference in diploid number and chromosome morphology between *Cebuella* and *Callithrix* is due to a Robertsonian-type fusion of chromosomes 16 and 22, and a paracentric inversion in chromosome 19 (Nagamachi et al, 1992).

*Geographical distribution.* *Cebuella* species range northwest Amazonia and are limited by the Madeira River in the south and by the Japurá River in the north; the southernmost records are in Ponton, Bolivia, and in the Manu National Park, Peru; the westernmost record is Copataza River, Equator, and the northernmost record is Puerto Limón, Putumayo, Colombia (Boubli et al, 2018).

*Evolutionary history.* The *Cebuella* clade branched off the *Mico*+*Callibella* ancestor during the upper Pliocene, around 3.41Mya, in the northwest Amazonia with the species splits occurring during the lower Quaternary, around 1.45Mya.

*Ecology.* Groups range in size from 2-9 individuals ($\overline{x}$=5.1) that inhabit edges and interiors of seasonally floodplain forests, edges of clearings, orchards, and primary and secondary forests, where they maintain small home ranges (0.1-0.5ha) determined by the spatial distribution of exudate source trees and vines they feed on in addition to arthropods, the main components of their diets (Soini, 1993).

*Etymology.* Derived from the Greek word kébos, monkey, and Latin diminutive suffix ellus (Barnett, 2004).

*Vernacular names.* sagui-pigmeu, leãozinho (Português), leoncillo, mono de bolsillo (Español), Pygmy marmoset (English), Gelbbauch-Zwergseidenäffchen (Deutsch), Ouistiti pygmée (Français).
